# Engineered probiotics limit CNS autoimmunity by stabilizing HIF-1α in dendritic cells

**DOI:** 10.1101/2023.03.17.532101

**Authors:** Liliana M. Sanmarco, Joseph M. Rone, Carolina M. Polonio, Federico Giovannoni, Gonzalo Fernandez Lahore, Kylynne Ferrara, Cristina Gutierrez-Vazquez, Ning Li, Anna Sokolovska, Agustin Plasencia, Camilo Faust Akl, Payal Nanda, Evelin S. Heck, Zhaorong Li, Hong-Gyun Lee, Chun-Cheih Chao, Claudia M. Rejano-Gordillo, Pedro H. Fonseca-Castro, Tomer Illouz, Mathias Linnerbauer, Jessica E. Kenison, Rocky M. Barilla, Daniel Farrenkopf, Gavin Piester, Lucas Dailey, Vijay K. Kuchroo, David Hava, Michael A. Wheeler, Clary Clish, Roni Nowarski, Eduardo Balsa, Jose M. Lora, Francisco J. Quintana

## Abstract

Dendritic cells (DCs) control the generation of self-reactive pathogenic T cells. Thus, DCs are considered attractive therapeutic targets for autoimmune diseases. Using single-cell and bulk transcriptional and metabolic analyses in combination with cell-specific gene perturbation studies we identified a negative feedback regulatory pathway that operates in DCs to limit immunopathology. Specifically, we found that lactate, produced by activated DCs and other immune cells, boosts NDUFA4L2 expression through a mechanism mediated by HIF-1α. NDUFA4L2 limits the production of mitochondrial reactive oxygen species that activate XBP1-driven transcriptional modules in DCs involved in the control of pathogenic autoimmune T cells. Moreover, we engineered a probiotic that produces lactate and suppresses T-cell autoimmunity in the central nervous system via the activation of HIF-1α/NDUFA4L2 signaling in DCs. In summary, we identified an immunometabolic pathway that regulates DC function, and developed a synthetic probiotic for its therapeutic activation.

## Introduction

Dendritic cells (DCs) are professional antigen-presenting cells with central roles in the control of adaptive immunity^1, 2^. In the context of autoimmune diseases, DCs regulate tissue pathology through a myriad of mechanisms, including the priming and differentiation of pathogenic and regulatory T cells^1, 2^, the local reactivation of autoimmune T cells in target tissues^3–5^, the control of epitope spreading^3–5^, and the regulation of microbiome-driven mechanisms of disease pathology^6^. Unsurprisingly, genetic variants associated with DC function have been linked to autoimmune disorders^7–10^, highlighting the central role of DCs in immune regulation. Based on these findings, DCs are considered attractive therapeutic targets for the management of human autoimmune diseases^11^. However, the design of successful therapies for autoimmunity requires the identification of regulatory mechanisms that control DC responses. Important mechanisms linked to DC regulation in cancer have been identified^12–14^, but less is known about their regulation in autoimmunity.

Metabolism regulates adaptive and innate immunity^15–17^. Indeed, the metabolism of DCs, T cells and other cell types controls autoimmune pathology in multiple setups. In addition, both host^18^ and commensal flora^19^ metabolites can regulate DC function, suggesting that probiotic-based approaches may allow the therapeutic modulation of DC responses in autoimmune diseases. However, un-manipulated probiotics use built-in anti-inflammatory mechanisms, evolutionarily selected to optimize host-commensal interactions in health but not in the context of autoimmune pathology^20^. In addition, the mechanism of action of un-manipulated probiotics is often unclear, limiting their utility for mechanistic studies. For example, *L. reuteri* is frequently used as a source of anti-inflammatory agonists for the aryl hydrocarbon receptor (AHR), but its therapeutic effects on neurologic disorders involve additional metabolites acting through AHR-independent pathways^21^. Synthetic biology allows the design of probiotics optimized for immune modulation via well-defined mechanisms of action, as exemplified by recent reports of engineered probiotics targeting L-arginine^22^ and purinergic signaling^23^ in the context of cancer immunotherapy and inflammatory bowel disease, respectively.

In this study, we combined single cell and bulk transcriptional and metabolic analyses with gene perturbation studies to investigate the regulation of DCs in autoimmunity. We found that L-lactate upregulates the expression of the NADH dehydrogenase (ubiquinone)-1α subcomplex 4-like 2 (NDUFA4L2) through a mechanism mediated by the transcription factor hypoxia inducible factor 1α (HIF-1α). NDUFA4L2 controls mitochondrial activity in DCs, limiting the expression of mitochondrial reactive oxygen species (mtROS)-driven transcriptional programs that promote the differentiation of encephalitogenic T cells. Moreover, we developed a synthetic probiotic engineered to produce lactate, which limits T cell-driven CNS autoimmunity in EAE via the activation of HIF-1α/NDUFA4L2 signaling in DCs. In summary, we identified a novel immunometabolic pathway which limits DC pro-inflammatory responses, and we engineered synthetic probiotics for its therapeutic modulation in autoimmune disorders.

### HIF-1α expression in DCs limits T-cell autoimmunity during EAE

To identify mechanisms involved in the regulation of DCs in CNS autoimmunity, we analyzed by scRNA-seq DCs recruited to the CNS in EAE induced in B6 mice by immunization with MOG_35-55_. We detected the activation of transcriptional programs associated with inflammation, such as NF-κB, PI3K/AKT and TLR signaling **(Fig. 1a)**. We also detected activation of signaling by the transcription factor HIF-1α in both type 1 and 2 classic DCs (cDC1s and cDC2s), where HIF-1α protein expression was also. **(Figs. 1a,b and Extended Data Figs. 1a-d)**. Indeed, we detected the upregulation of HIF-1α protein levels in CNS, lymph nodes and spleen DCs at the onset of EAE **(Figs. 1c,d)**.

**Figure 1.**
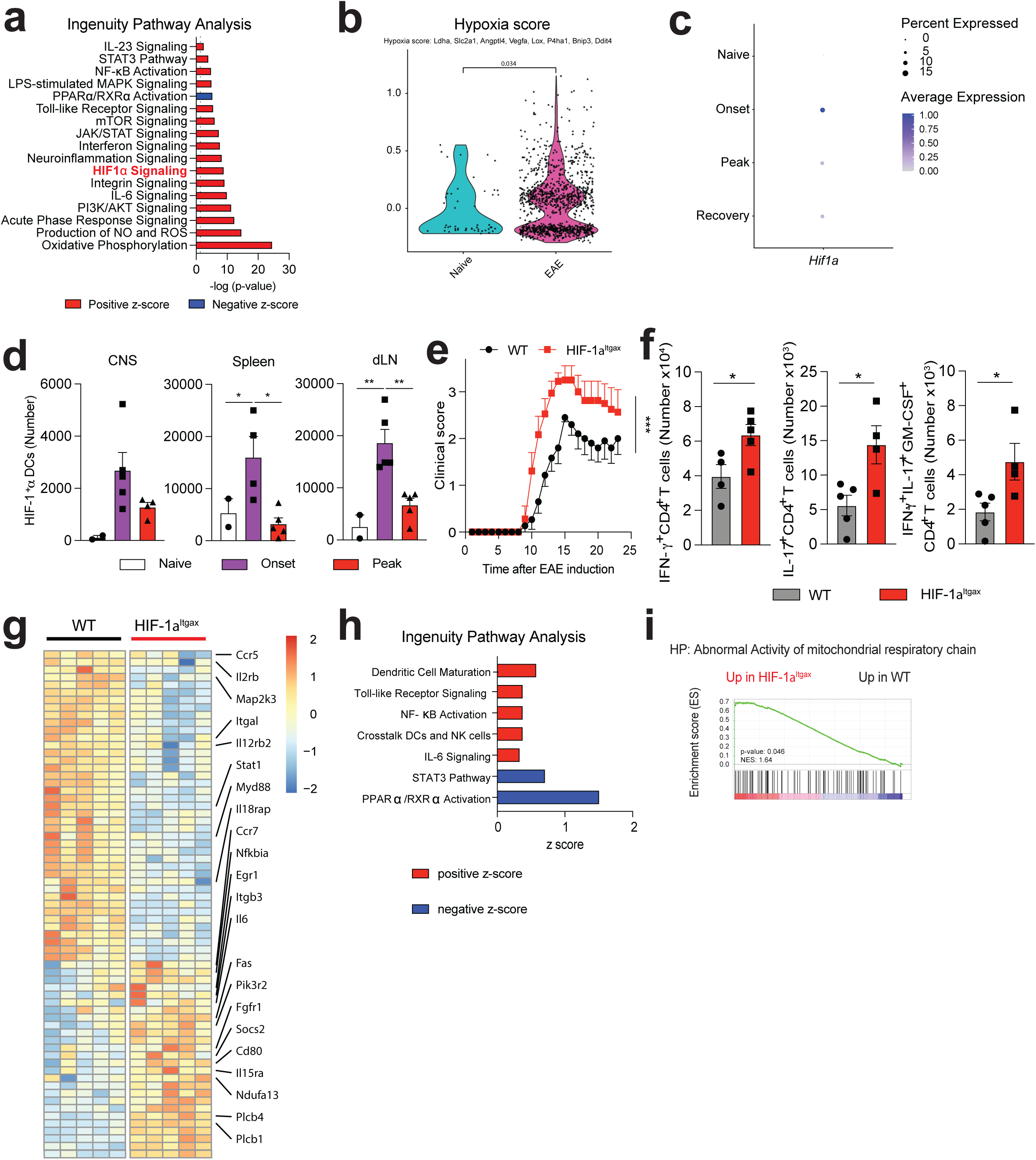
HIF-1α in DCs limits T-cell autoimmunity during EAE. **(a-c)** Ingenuity pathway analysis (IPA) (a), hypoxia score (b) and *Hif1a* expression (c) in total DCs from the CNS of EAE mice. **(d)** HIF-1α expression in DCs isolated from CNS, lymph nodes and spleen of WT mice before (“naive” n=2), 12 (“onset” n=5) or 17 days after EAE induction (“peak” n=5). **(e)** EAE in WT (n=15) and HIF-1α^Itgax^ (n=9) mice. Experiment repeated three times. **(f)** IFN-γ^+^, IL-17^+^, and IFN-γ^+^IL-17^+^GM-CSF^+^ CD4^+^ T cells in the CNS of WT and HIF-1α^Itgax^ mice 15 days after EAE induction. n=5 mice per group. Unpaired t-test. **(g)** RNA-seq analysis of splenic DCs at peak EAE. n=5 mice per group. **(h,i)** IPA (h) and Gene Set Enrichment Analysis (GSEA) (i) in WT and HIF-1α^Itgax^ splenic DCs at peak EAE. Data shown as mean±SEM. ****p<0.0001, ***p<0.001, **p<0.01, *p<0.05, ns: p>0.05.

HIF-1α-driven transcriptional responses have been studied in the context of tumor immune evasion^24^, but much less is known about the control of DCs by HIF-1α during autoimmunity. To study the role of HIF-1α in DCs during CNS inflammation we generated Itgax^Cre^HIF-1α^flox^ (HIF-1α^Itgax^) mice. HIF-1α deletion in DCs resulted in the worsening of EAE, concomitant with increased IFN-γ^+^, IL-17^+^ and GM-CSF^+^ CD4^+^ T-cell numbers in the CNS and the periphery, and increased recall responses to MOG_35-55_ **(Figs. 1e,f and Extended Data Figs. 1e-g)**; DC recruitment to the CNS was unaffected **(Extended Data Fig. 1i)**.

RNA-seq analyses of HIF-1α-deficient splenic DCs detected increased activation of pathways linked to inflammation and abnormal mitochondrial respiratory function during EAE **(Figs. 1g-i)**. Similarly, HIF-1α-deficient DCs isolated from the CNS during EAE showed increased transcriptional responses linked to neuroinflammation, ROS production, and response to mitochondrial depolarization **(Extended Data Figs. 1j-l, Table 1)**.

**Table 1.** Genes associated to GSEA (GO:0051882) response to mitochondria depolarization.

To validate these findings, we analyzed the response of bone marrow-derived DCs (BMDCs) to hypoxia (1% oxygen) and other stimuli that stabilize HIF-1α and increase its intracellular levels such as deferoxamine^25^, ML228^26^ and the knockdown of the Von Hippel– Lindau tumor suppressor (VHL)^25^. HIF-1α stabilization decreased *Il1b*, *Il6*, *Il23*, *Tnf* and *Il12* expression in DCs **(Fig. 2a, Extended Data Fig. 2a,b),** and *Ifng* and *Il17* expression in co-cultured 2D2^+^ CD4^+^ T cells in the presence of their cognate antigen MOG35-55 **(Extended Data Fig. 2c)**. HIF-1α stabilization did not affect DC viability **(Extended Data Figs. 2d,e)**. To further investigate the role of HIF-1α in the control of the T-cell response, we infected HIF-1α^Itgax^ mice with *Salmonella typhimurium* expressing the T-cell epitope 2W1S^27^ **(Extended Data Fig. 2f)**. HIF-1α deletion in DCs resulted in lower cecal bacteria burden concomitant with an increase in 2WS1-specific CD4^+^ T cells, and IFN-γ^+^ CD4^+^ and IL-17^+^ CD4^+^ T cells in the colon **(Extended Data Figs. 2g-i)**. Taken together, these findings suggest that HIF-1α limits DC transcriptional programs that promote effector T-cell responses.

**Figure 2.**
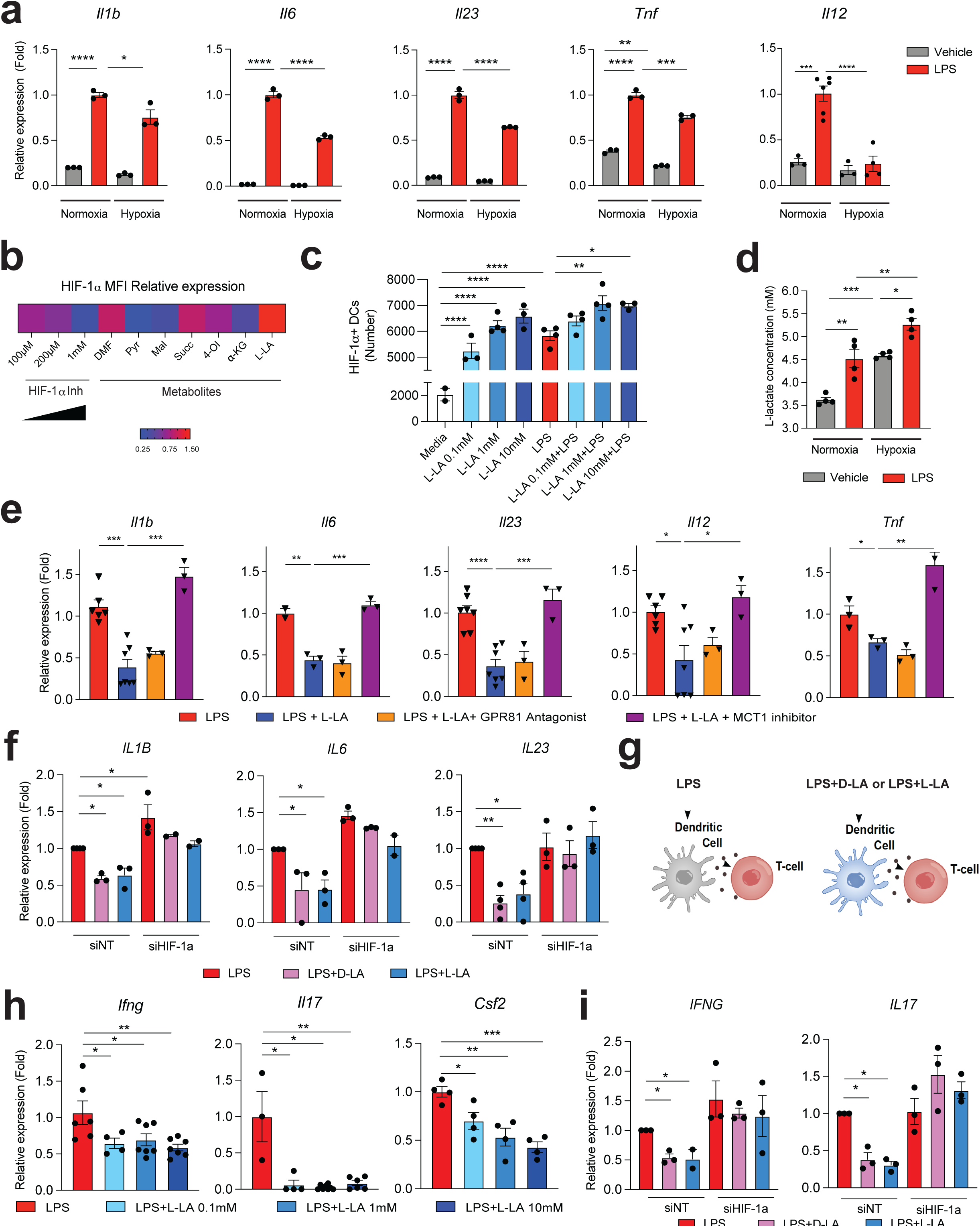
HIF-1α activation by lactate inhibits DC pro-inflammatory activities. **(a)** mRNA expression in DC activated with LPS under normoxic or hypoxic conditions for 6 h. n=3 per group. **(b)** Mean fluorescence intensity (MFI) of HIF-1α expression in splenic DCs after metabolite treatment and LPS activation for 6h, normalized to LPS stimulation. n=4 per condition. DMF: dimethyl fumarate, Pyr: pyruvate, Mal: Malonate, Succ: Succinate, 4-OI: 4-octyl itaconate, αKG: αketoglutarate, L-LA: L-sodium lactate. **(c)** HIF-1α^+^ DCs following treatment with L-LA (0.1, 1 and 10 mM) and LPS for 6h. **(d)** L-LA levels following DC activation with LPS under normoxia or hypoxia. **(e)** mRNA expression in WT BMDCs after treatment with LPS, L-LA, GPR81 antagonist or MCT1 inhibitor. **(f)** mRNA expression in human DCs stimulated with LPS, D-LA or L-LA (1mM) after HIF-1α knockdown. **(g,h)** Experimental design (g) and *Ifng*, *Il17* and *Csf2* expression (h) in 2D2^+^ CD4^+^ T cells co-cultured with WT BMDCs pre-stimulated with L-LA and LPS. **(i)** *IFNG* and *IL17* expression in human T cells co-cultured with DC from Fig. 2f. Data shown as mean±SEM. ****p<0.0001, ***p<0.001, **p<0.01, *p<0.05, ns: p>0.05.

### HIF-1α activation by lactate inhibits DC pro-inflammatory responses

Metabolites can stabilize HIF-1α^28^. Thus, we evaluated the effect of metabolites on HIF-1α intracellular levels in splenic DCs. L-lactate (L-LA) induced the highest expression of HIF-1α **(Figs. 2b,c, Extended Data Fig. 3a,b)**, and activated HIF-1α-driven gene expression in BMDCs from ODD-Luc mice, which harbor a HIF-1α-responsive luciferase reporter^29^ **(Extended Data Fig. 3c)**. Interestingly, LPS increased L-LA production, *Hif1a* expression and HIF-1α levels in DCs **(Figs. 2c,d, Extended Data Fig. 3d)**, suggesting that DC activation induces both *Hif1a* expression and lactate-dependent HIF-1α stabilization.

**Figure 3.**
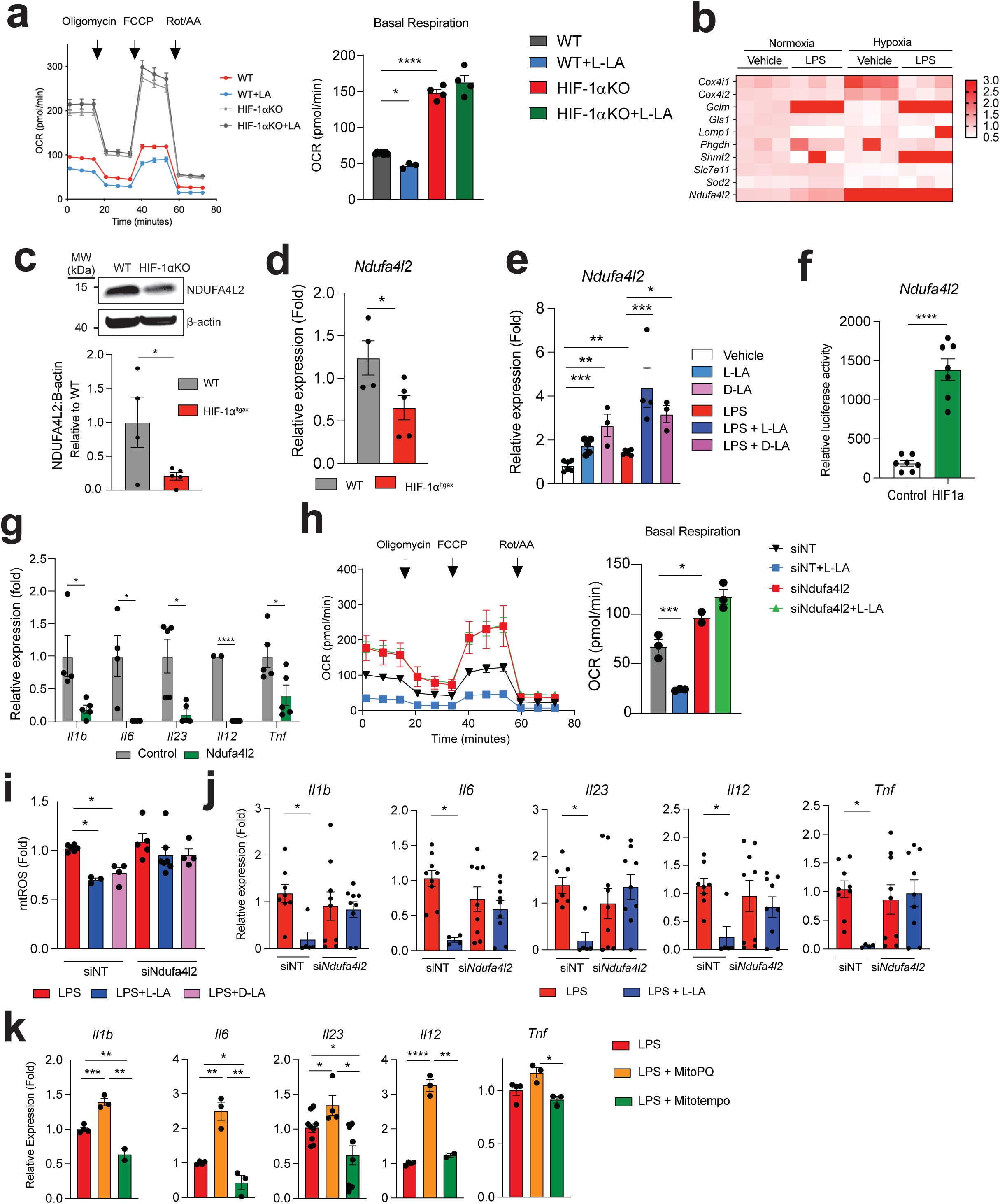
HIF-1α-induced NDUFA4L2 limits mtROS production. **(a)** Oxygen consumption rate (OCR) in WT and HIF-1α^Itgax^ BMDCs after L-LA stimulation. **(b)** Expression of HIF-1α-target genes in BMDCs stimulated with LPS under normoxic or hypoxic conditions. **(c)** Western blot of NDUFA4L2 expression in WT and HIF-1α^Itgax^ BMDCs after LPS stimulation. **(d)** *Ndufa4l2* expression in WT and HIF-1α^Itgax^ splenic DCs at EAE peak. **(e)** *Ndufa4l2* expression after 6h treatment with L-LA, D-LA or LPS. **(f)** Transactivation of *Ndufa4l2* promoter following HIF-1α over-expression. **(g)** Effect of *Ndufa4l2 overexpression in* LPS-stimulated BMDCs. **(h-j)** Effect of *Ndufal2* knockdown on OCR (h), mtROS production (i) and gene expression (j) in BMDCs treated with LPS, L-LA or D-LA. **(k)** mRNA expression in BMDCs treated with LPS, MitoParaquat (MitoPQ) or MitoTempo for 6h. Data shown as mean±SEM. ***p<0.001, **p<0.01, *p<0.05, ns: p>0.05.

L-LA treatment decreased pro-inflammatory cytokine expression induced by LPS in primary murine and human DCs **(Figs. 2e,f)**. In addition, L-LA pre-treatment of DCs decreased *Ifng*, *Il17* and *Csf2* expression in 2D2^+^ CD4^+^ T cells co-cultured in the presence of MOG_35-55_ **(Figs. 2g,h)**. These anti-inflammatory effects of L-LA where dose-dependent, and involved the monocarboxylate transporter (MCT) 1, which transports L-LA across the cell membrane^30^, but not the GPR81 membrane receptor^31^ **(Figs. 2e,h, Extended Data Figs. 3e-h)**. Moreover, the anti-inflammatory effects of L-LA on DCs were also HIF-1α-dependent in mice and human DCs **(Figs. 2f,i, Extended Data Figs. 4a-c)**. These findings suggest that DC activation triggers an L-LA/HIF-1α-driven anti-inflammatory feedback loop.

**Figure 4.**
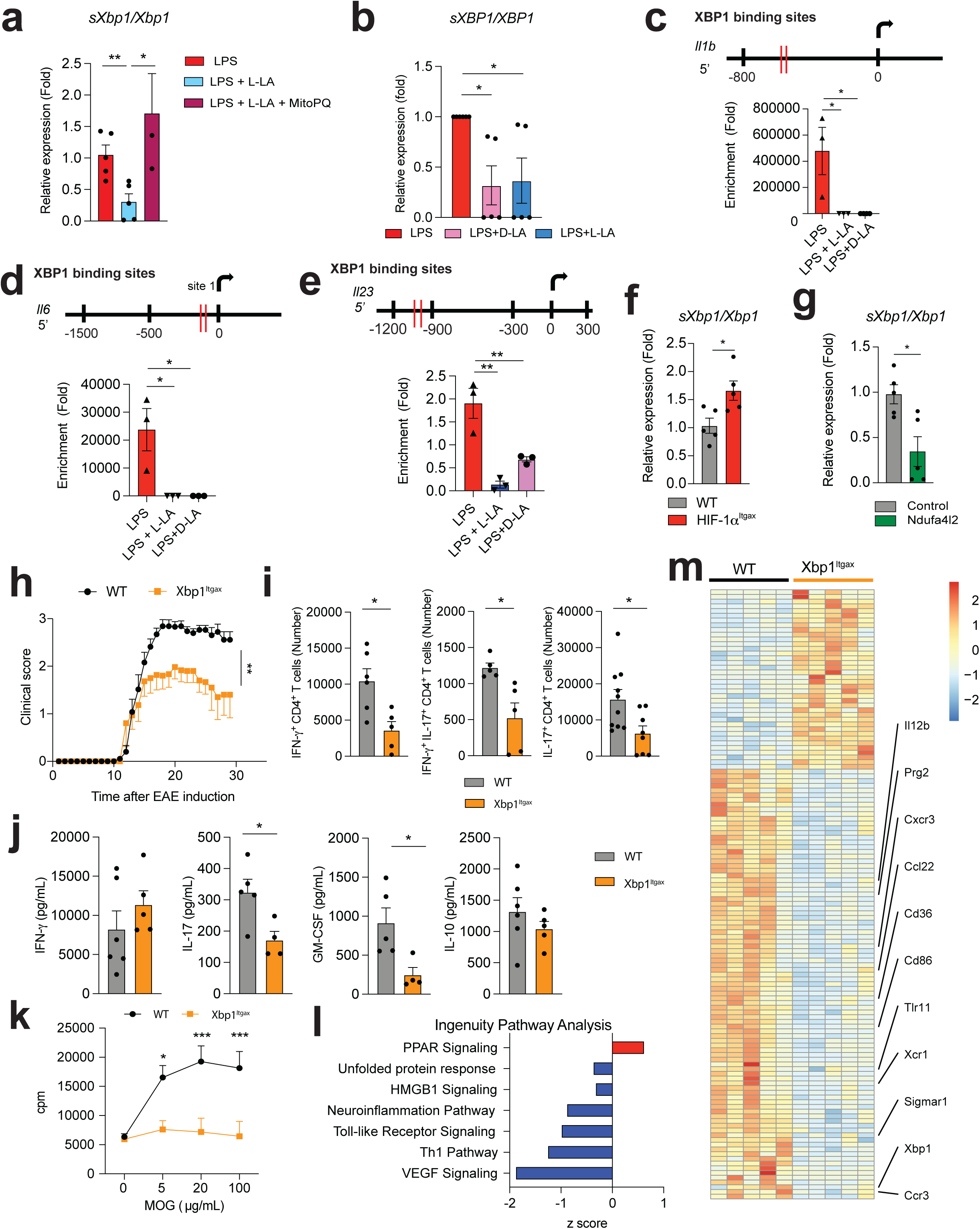
L-LA limits mtROS-driven activation of XBP1 in DCs. **(a)** *Spliced Xbp1 mRNA* normalized to *total Xbp1 mRNA (sXbp1/Xbp1)* in BMDCs treated with LPS, L-LA or MitoPQ for 6h. **(b)** *sXPB1/XBP1* in human DCs treated with LPS, D-LA or L-LA for 6h. **(c-e)** XBP1 biding sites and XBP1 recruitment to *Il1b* (c), *Il6* (d) and *Il23* (e) promoters in BMDCs determined by ChIP after treatment with LPS, D-LA or L-LA for 6h. **(f)** *sXbp1/Xbp1* in WT and HIF-1α^Itgax^ splenic DCs at EAE peak. **(g)** *sXbp1/Xbp1* in BMDCs over-expressing *Ndufa4l2*. **(h)** EAE in WT (n=15) and XBP1^Itgax^ (n=15) mice. Experiment repeated three times. **(i)** IFN-γ^+^, IFN-γ^+^IL-17^+^, and IL-17^+^ CD4^+^ T cells in CNS. **(j,k)** Recall cytokine (j) and proliferative (k) response following stimulation of WT and Xbp1^Itgax^ splenocytes with 20 μg/mL of MOG35-55 for 72h. n=5 mice per group. **(l,m)** IPA (l) and heatmap (m) of RNA-seq analysis of WT and Xbp1^Itgax^ splenic DCs 30 days after EAE induction. n=5 mice per group. Data shown as mean±SEM. ***p<0.001, **p<0.01, *p<0.05, ns: p>0.05.

Two lactate stereoisomers exist in nature: L-LA is produced by mammalian cells, while D-lactate (D-LA) is provided by the diet, the microbiome and the methylglyoxal pathway^32^. D-LA and L-LA induced similar HIF-1α protein upregulation in mouse DCs **(Fig. 2c, Extended Data Fig. 4d)**. Moreover, D-LA recapitulated L-LA anti-inflammatory activities, decreasing pro-inflammatory cytokine expression in mouse and human DCs and reducing effector T-cell polarization in co-culture studies **(Figs. 2f,i, Extended Data Figs. 4e,f)**. Thus, HIF-1α activation by L-LA or D-LA limits pro-inflammatory DC responses.

### HIF-1α-induced NDUFA4L2 limits mtROS production

HIF-1α controls multiple biological processes including metabolism^33^. Indeed, HIF-1α-deficient DCs showed decreased glucose transporter expression, glycolytic activity **(Extended Data Figs. 5a-c)**. Moreover, HIF-1α-deficiency resulted in increased oxygen consumption rate (OCR); L-LA decreased the OCR in WT but not in HIF-1α^Itgax^ DCs **(Fig. 3a)**. Based on the detection of transcriptional modules linked to abnormal activity of mitochondrial respiratory chain in HIF-1α^Itgax^ DCs **(Fig. 1i)** and the HIF-1α-dependent effects of L-LA on the OCR **(Fig. 3a)**, we analyzed the expression of mitochondrial HIF-1α-target genes in BMDCs. We detected a HIF-1α-dependent increase in *Ndufa4l2* expression following DC stimulation or treatment with L-LA or D-LA **(Figs. 3b-e)**. Moreover, LPS, L-LA, D-LA and HIF-1α overexpression transactivated the *Ndufa4l2* promoter in reporter assays **(Fig. 3f and Extended Data Fig. 5d)**.

**Figure 5.**
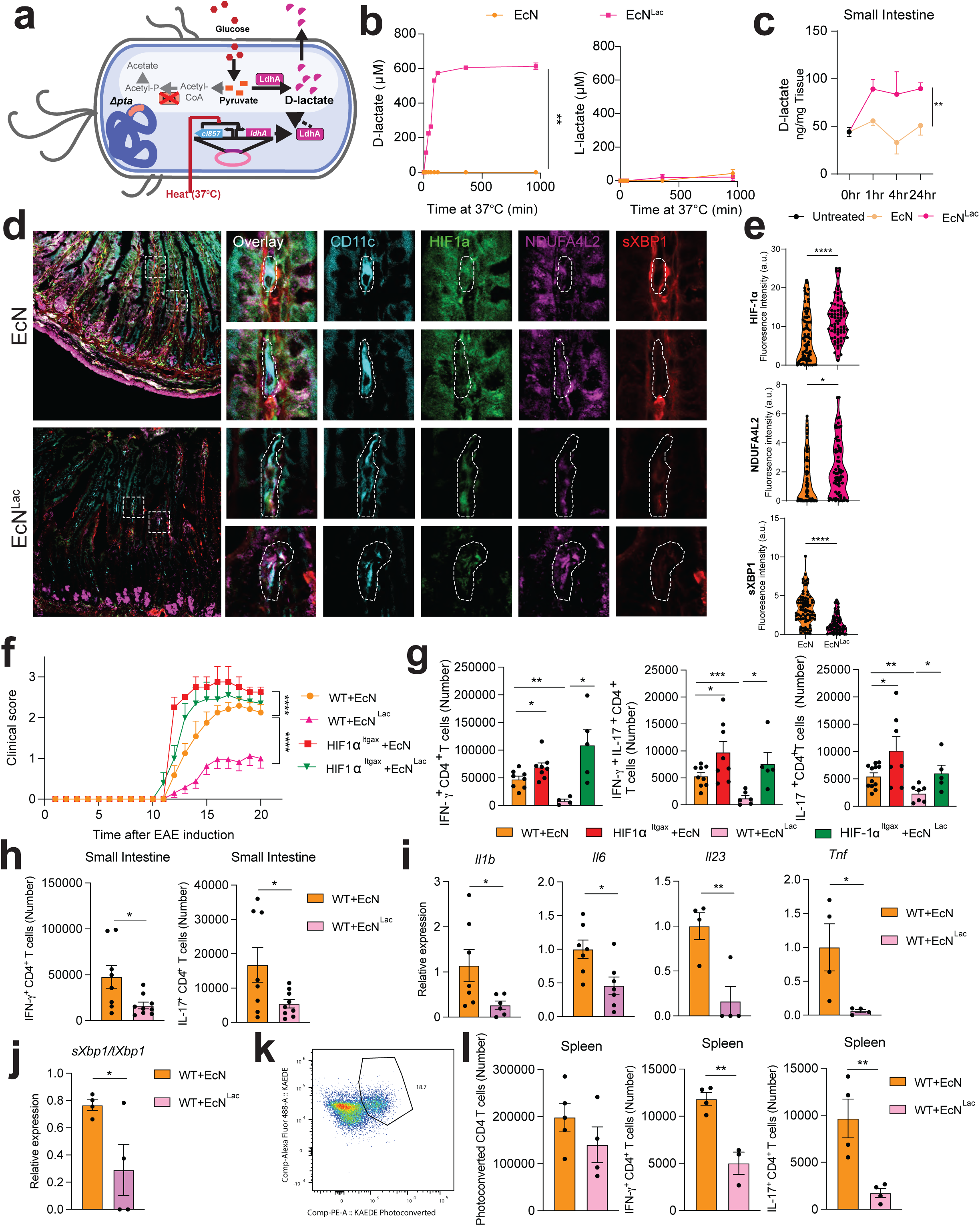
HIF-1α/NDUFA4L2 activation by engineered probiotics ameliorates EAE. **(a)** Engineered EcN^Lac^ strain. **(b)** D-LA and L-LA in culture supernatants following EcN^Lac^ or EcN incubation at 37°C. **(c)** D-LA in small intestine tissue after EcN^Lac^ administration. **(d,e)** HIF-1α, NDUFA4L2 and spliced XBP1 protein staining (d) and quantification (e) in CD11c^+^ cells from small intestine after EcN^Lac^ or EcN gavage. **(f)** EAE development and **(g)** IFN-γ^+^, IL-17^+^ and IFN-γ^+^ IL-17^+^ CD4^+^ T cell quantification in CNS of WT and HIF-1α^Itgax^ mice treated with EcN or EcN^Lac^. Experiment repeated three times. **(h)** IFN-γ^+^ and IL-17^+^ CD4^+^ T cell quantification in small intestine of WT mice treated with EcN or EcN^Lac^. (**i,j)** *Il1b, Il6, Il23, Tnf* (i) and *sXbp1/Xbp1* (j) expression in small intestine DCs from EcN or EcN^Lac^ treated mice. **(k)** Representative dot plot of Kaede photoconversion in T cells from spleen. **(l)** Total, IFN-γ^+^ and IL-17^+^ CD4^+^ T cell photoconverted in spleen of WT mice treated with EcN or EcN^Lac^. Data shown as mean±SEM. **p<0.01, *p<0.05, ns: p>0.05.

NDUFA4L2 is a subunit of respiratory Complex I which limits its activity and the generation of mitochondrial reactive oxygen species (mtROS)^34^. *Ndufa4l2* overexpression abrogated the increase in basal respiration and proton leakage induced by HIF-1α deletion, suggesting that HIF-1α-driven *Ndufa4l2* expression limits mitochondrial respiration in DCs **(Extended Data Figs. 5e,f)**. Moreover, based on the reported control of innate and adaptive immune cells by mitochondria^35^, we analyzed the role of *Ndufa4l2* on DC responses. *Ndufa4l2* overexpression limited pro-inflammatory cytokine expression in DCs **(Fig. 3g)**.

mtROS is a byproduct of Complex I activity with pro-inflammatory properties^36^. L-LA and D-LA limited mtROS production and oxygen consumption in DCs in an *Ndufa4l2*-dependent manner, as *Ndufa4l2* knockdown suppressed the effects of L-LA on the OCR and mtROS production **(Figs. 3h,i, Extended Data Fig. 5g)**. In addition, *Ndufa4l2* knockdown abrogated the anti-inflammatory effects of L-LA and D-LA on DCs **(Fig. 3j)**.

To investigate the link between lactate, mtROS and DC responses, we used mitoParaquant (mitoPQ), which selectively increases mtROS production, and the mtROS scavenger mitoTempo. MitoPQ boosted LPS-induced DC pro-inflammatory gene expression, while mitoTempo suppressed it **(Fig. 3k)**. Moreover, mitoPQ abolished the inhibitory effects of L-LA on the expression of pro-inflammatory cytokines by DCs **(Extended Data Figs. 6a,b)**, suggesting that the NDUFA4L2-dependent suppression of mtROS production mediates the anti-inflammatory effects of lactate in DCs.

L**-**LA can be converted to pyruvate to enter the tricarboxylic acid (TCA) cycle^37^. However, using uniformly labeled 13C lactate we detected very low label incorporation (about 1%) into TCA intermediates in BMDCs **(Extended Data Fig. 6c)**. In addition, L-LA or D-LA treatment did not increase intracellular pyruvate levels **(Extended Data Fig. 6d)**, and inhibition of L-LA oxidation into pyruvate by oxamate did not abrogate LA tolerogenic activities on DCs **(Extended Data Figs. 6e,f)**, overall suggesting that LA conversion into pyruvate and its use as fuel does not contribute to LA anti-inflammatory effects in DCs. Indeed, pyruvate did not suppress DC pro-inflammatory gene expression **(Extended Data Fig. 6g)**. Taken together, these data suggest that by limiting Complex I activity, LA/HIF-1α-driven NDUFA4L2 expression limits mtROS-induced pro-inflammatory DC responses.

### L-LA limits mtROS-driven activation of XBP1 in DCs

ROS^36^ and inflammation^38^ activate the unfolded protein response (UPR) and the transcription factor XBP1, which drives pro-inflammatory responses in DCs^36^ and macrophages^38^. Specifically, ROS^36^ and inflammation^38^ trigger the excision of a intron from the *Xbp1*mRNA, generating a spliced *Xbp1* mRNA (sXbp1), which codes for a full-length functional XBP1^39^. L-LA reduced XBP1 activation in mouse BMDCs and human DCs, as indicated by the analysis of *Xbp1* mRNA splicing, but this effect was lost when mtROS generation was induced with mitoPQ **(Figs. 4a,b, Extended Data Fig. 7a,b)**. Moreover, L-LA reduced the recruitment of XBP1 to the *Il1b*, *Il6* and *Il23* promoters triggered by LPS **(Figs. 4c-e)**. Conversely, mtROS production triggered by mitoPQ boosted LPS-induced XBP1 recruitment to pro-inflammatory cytokine promoters **(Extended Data Fig. 7c)**.

In support for a role of HIF-1α in limiting XBP1 activation, we detected increased *Xbp1* mRNA splicing in HIF-1α-deficient splenic DCs during EAE **(Fig. 4f)**, and reduced XBP1 recruitment to pro-inflammatory cytokine promoters after HIF-1α stabilization with ML-228 **(Extended Data Fig. 7d)**. Similarly, *Ndufa4l2* over-expression reduced LPS-induced *Xbp1* mRNA splicing **(Fig. 4g)**, suggesting that HIF-1α-driven *Ndufa4l2* expression limits XBP1 activation.

Finally, we induced EAE on Itgax^Cre^Xbp1^flox^ (Xbp1^Itgax^) mice^40^. EAE was ameliorated in Xbp1^Itgax^ mice, concomitant with a reduction in CNS and splenic IFNγ^+^, IL-17^+^ and IFNγ^+^ IL-17^+^ CD4^+^ T cells, and a decreased recall response to MOG35-55 **(Figs. 4h-k, Extended Data Figs. 7e)**. In RNA-seq analyses, XBP1-deficient DCs displayed a decreased pro-inflammatory response **(Figs. 4l,m, Extended Data Figs. 7f,g)**. Collectively, these findings suggest that by limiting mtROS production, L-LA/HIF-1α-driven *Ndufa4l2* expression suppresses XBP1-dependent pro-inflammatory DC responses.

### HIF-1α/NDUFA4L2 activation by engineered probiotics ameliorates EAE

Finally, we studied the therapeutic potential of activating HIF-1α/NDUFA4L2 signaling. L-LA or D-LA administration (125 mg/kg, daily) suppressed the development of encephalitogenic T cells and EAE induced by MOG_35-55_ immunization **(Extended Data Figs. 8a,b)**, and limited gene expression linked to inflammation mitochondrial metabolism, oxidative stress and the UPR in DCs **(Extended Data Figs. 8c-f)**. Of note, D-LA and L-LA plasma levels were similar in EAE and naïve mice **(Extended Data Fig. 9a)**.

To facilitate the intestinal absorption of lactate following oral delivery while minimizing the potential for lactic acidosis associated with bolus lactate administration^32^, we engineered probiotics that produce D-LA using as a chassis the *E. coli Nissle* strain (EcN^Lac^), which has already been used to develop probiotics tested in humans^22, 41, 42^ **(Fig. 5a)**. Specifically, we knocked out the *pta* gene on *E. coli Nissle* (EcN), and introduced a plasmid harboring the *ldhA* gene under the control of a heat inducible promoter active at 37°C to ensure sufficient conversion of pyruvate to D-LA **(Fig. 5a, Extended Data Figs. 9b,c)**^43, 44^. Sequencing of the bacterial chromosome confirmed *pta* deletion **(Extended Data Fig. 9b).**

We first evaluated the ability of the heat inducible promoter to drive green fluorescent protein (GFP) expression in an EcN strain (EcN^GFP^), detecting GFP expression 20min after heat activation at 37°C **(Extended Data Figs. 9d,e)**. Similarly, the EcN^Lac^ strain, but not the parental EcN strain, secreted D-LA when cultured at 37°C **(Fig. 5b)**; we did not detect L-LA production by EcN^Lac^ **(Fig. 5b)**.

To characterize the distribution and biologic effects of probiotic-produced D-LA, we administered EcN^Lac^ or EcN by oral gavage daily for a week, and quantified D-LA levels in plasma, small intestine and colon 1, 4 and 24 h after the last administration of the probiotic. EcN^Lac^ administration increased D-LA levels in plasma, and also in small intestine and colon tissue, but not in cerebrospinal fluid (CSF) or CNS lysates **(Fig. 5c, Extended Data Figs. 9f-h)**. Moreover, EcN^Lac^ gavage increased HIF-1α and NDUFA4L2 protein levels and HIF-1α signaling in small intestine DCs, while decreasing the expression of active XBP1 protein **(Figs. 5d,e Extended Data Figs. 9i-k)**. Of note, neither EcN^Lac^ nor EcN^GFP^ were detected in the bloodstream after oral gavage **(Extended Data Figs. 9l,m)**. Taken together, these findings suggest that the local increase in D-LA resulting from EcN^Lac^ administration activates HIF-1α/NDUFA4L2 signaling in intestinal DCs.

To evaluate the effects on CNS inflammation of HIF-1α/NDUFA4L2 activation, we administered EcN^Lac^ or EcN daily or weekly by gavage starting 3 days before EAE induction **(Extended Data Fig. 9n)**. EcN^Lac^ administration ameliorated EAE, decreasing pro-inflammatory T cells in the CNS, small intestine and periphery and the recall response to MOG_35-55_ **(Figs. 5f-h, Extended Data Figs. 10a-g)**. However, EcN^Lac^ administration, did not affect the bacterial burden nor pathogen-specific CD4^+^ T cells following challenge with *S. typhimurium* **(Extended Data Figs. 10h-k)**.

The intestinal microbiome regulates pathogenic T cells in EAE^45^. Indeed, recent studies documented the priming in intestinal tissues of encephalitogenic T cells which then migrate to the spleen and CNS during EAE^46, 47^. Thus, we analyzed the effects EcN^Lac^ administration on intestinal DCs and T cells. EcN^Lac^ administration increased HIF-1α intracellular levels DCs but not in neutrophils, monocytes or T cells in the small intestine **(Fig. 5d,e, Extended Data Fig. 10l,m)**. Indeed, the reduction of pathogenic T-cell responses and the amelioration of EAE by EcN^Lac^ were abrogated in HIF-1α^Itgax^ mice, supporting a role for HIF-1α expressed in DCs in the anti-inflammatory effects of EcN^Lac^ **(Figs. 5f,g and Extended Data Figs. 10a)**. In agreement with this interpretation, intestinal DCs from EcN^Lac^-treated mice displayed reduced *Xbp1* mRNA splicing and decreased expression of pro-inflammatory cytokines **(Figs. 5i,j)**. Moreover, we detected a reduction in IFNγ^+^ CD4^+^ and IL-17^+^ CD4^+^ T cells that migrated from the intestine to the spleen as determined by the use of *CAG^Kaede^*mice expressing a photoconvertible fluorescent protein^48^ **(Figs. 5k,l)**; the total number of CD4^+^ T cells migrating from the intestine to the spleen remained unaffected **(Fig. 5l)**. These findings suggest that EcN^Lac^ suppress CNS inflammation via the D-LA-driven activation of HIF-1α/NDUFA4L2 in intestinal DCs and, potentially, other cell types.

## Discussion

Immunosuppressive roles of HIF-1α in the tumor microenvironment have reported, but less is known about its role in autoimmunity^33, 49, 50^. Hypoxia is considered the bona fide HIF-1α activator ^33^, but metabolites such as succinate^28^, fumarate^51^, oxaloacetate^52^, citrate^52^ and iron chelating agents^25^, can also activate HIF-1α. Here we report that lactate-triggered HIF-1α activation induces the expression of NDUFA4L2, which promotes mitochondrial fitness, limiting mtROS/XBP1-driven pro-inflammatory responses in DCs. DC stimulation by LPS or other microbial molecules promotes NF-κB-driven *Hif1a* expresssion^53^, increasing the levels of HIF-1α available for stabilization by lactate, and consequently limiting via NDUFA4L2 the mtROS-driven XBP1 activation of pro-inflammatory responses in DCs. Together with previous reports^17^, our findings suggest that an L-LA/HIF-1α/NDUFA4L2 signaling axis participates in a negative feedback loop which limits DC responses to minimize immunopathology, as described for IL-10^54^ and IL-27^55^. These findings are reminiscent of the Warburg effect^56^, the aerobic glycolysis and high production of L-LA associated with the promotion of metastasis, angiogenesis and tumor progression.

Lactate^57^ and HIF-1*α*^58^ have been linked to epigenetic and metabolic mechanisms of trained immunity in monocytes and macrophages. In addition, lactate is reported to boost the immunosuppressive activity of macrophages^24, 57^ and Foxp3^+^ Tregs^59^. Hence, additional mechanisms besides the activation of HIF-1α/NDUFA4L2 signaling in DCs may contribute to the suppression of EAE by EcN^Lac^. Indeed, the increased response to *S. typhimurium* infection of HIF-1α^Itgax^ mice suggest a general immune activation which may obscure the contribution of other cell types besides DCs to the suppressive effects of EcN^Lac^ on EAE.

The diet and the commensal microbiota are a physiologic source of D-LA, which may activate HIF-1α in DCs and other cells in intestinal tissue as part of homeostatic anti-inflammatory mechanisms, resembling recent observations made for AHR^60, 61^. Interestingly, both HIF-1α^24^ and AHR^62, 63^ are hyperactivated as part of tumor immunoevasion strategies, and HIF-1α and AHR transcriptional activity requires dimerization with the same cofactor, HIF-1β/ARNT^50, 64^. These findings suggest a potential crosstalk between HIF-1α and AHR signaling in DCs, as previously described in T cells^65^. Nevertheless, considering the paradoxical immunosuppressive effects of chronic XBP1 activation in the tumor microenvironment^40^, future studies should investigate the effects of chronic HIF-1α/NDUFA4L2 signaling on XBP1-driven transcriptional programs.

In summary, we identified a lactate-driven HIF-1α/NDUFA4L2 signaling axis which operates in DCs to limit XBP1-driven transcriptional modules that promote T-cell autoimmunity. Our studies also identify HIF-1α/NDUFA4L2 signaling as a candidate target for therapeutic intervention. Indeed, several immunomodulatory DC-targeting approaches have been developed, including nanoliposomes engineered to co-deliver autoantigen and a tolerogenic AHR agonist^66^, or to deliver an autoantigen-encoding modified mRNA^67^. Engineered probiotics provide an alternative approach for therapeutic immunomodulation, enabling the chronic administration of therapeutic agents with minimal adverse effects. Indeed, its established safety profile, increased serum sensitivity, broad antibiotic susceptibility, defined genomic landscape and high engineerability^68–70^ support the use of EcN for the therapeutic activation of HIF-1α/NDUFA4L2 signaling in autoimmunity. The development of probiotics engineered to activate immuno-regulatory signaling pathways may provide new tools for the clinical management of autoimmune and allergic disorders.

## Acknowledgments

This work was supported by grants NS102807, ES02530, ES029136, and AI126880 from the NIH; RG4111A1 and JF2161-A-5 from the National Multiple Sclerosis Society; RSG-14-198-01-LIB from the American Cancer Society; and PA-160408459 from the International Progressive MS Alliance (to F.J.Q.). F.J.Q. was supported by the DFG-funded CRC/TRR167 “NeuroMac.” C.M.P. was supported by a fellowship from FAPESP BEPE (2019/13731-0). C.G.-V. was supported by an Alfonso Martin Escudero Foundation postdoctoral fellowship and by a postdoctoral fellowship (ALTF 610-2017) from the European Molecular Biology Organization. G.F.L. received support by a grant from the Swedish Research Council (2021-06735). C.-C.C. received support from postdoctoral research abroad program (104-2917-I-564-024) from the Ministry of Science and Technology, Taiwan. C.M. R.-G. was supported by a predoctoral F.P.U. fellowship of the Ministry of Economy and Competitiveness and by the European Union FEDERER program. M.A.W. was supported by NIH (1K99NS114111, F32NS101790, T32CA207201), the Program in Interdisciplinary Neuroscience and the Women’s Brain Initiative at Brigham and Women’s Hospital. T.I. was supported by the EMBO postdoctoral fellowship (ALTF: 1009-2021). H.-G.L. was supported by a Basic Science Research Program through the National Research Foundation of Korea (NRF) funded by the Ministry of Education (2021R1A6A3A14039088). We thank Dr. Laurie Glimcher and Dr. Juan R. Cubillos Ruiz for sharing Itgax^Cre^Xbp1f^lox^ mice. We thank Dr. Stephen McSorley for providing the Salmonella typhimurium strain and Dr. Huixin Xu and Dr. Maria Lehtinen for training on CSF extraction. We thank all members of the Quintana laboratory for helpful advice and discussions; R. Krishnan for technical assistance with flow cytometry studies and the NeuroTechnology Studio at Brigham and Women’s Hospital for providing Seahorse instrument access.

## Author contributions

L.M.S., J.M.R., C.M.P., F.G., G.F.L., K.F., C.G.-V., N.L., A.S., A.P., C.F.A., P.N., E.S.H., H.G.L., C.-C.C., C.M.R.-G., P.H.F.-C., M.L., J.E.K.-W., R.M.B., D.F., G.P., E.N.C., and L.D. performed *in vitro* and *in vivo* experiments, FACS and genomic experiments. L.M.S., Z.L. and M.A.W. performed bioinformatic analyses. T.I. performed unbiased quantification of histology samples. J.M.R., V.K.K. and R.N. contributed reagents. L.M.S., N.L., D.H., A.S., J.M.L., F.J.Q. designed and generated engineered probiotics. L.M.S., E.B. and F.J.Q. discussed and interpreted findings and wrote the manuscript with input from coauthors. F.J.Q. designed and supervised the study and edited the manuscript.

## Materials and Methods

### Animals

Adult female mice were used on a C57Bl/6J background (The Jackson Laboratory, #000664) except for HIF-1α reporter FVB.129S6-Gt(ROSA)26Sortm2(HIF1A/luc)Kael/J mice (The Jackson Laboratory, #006206). B6.Cg-Tg(Itgax-cre)1-1Reiz/J mice (The Jackson Laboratory, #008068) were bred to B6.129-*Hif1a^tm3Rsjo^*/J mice (The Jackson Laboratory, #007561) to generate Itgax^Cre^HIF-1α^fl^ mice. Itgax^Cre^XBP1^fl^ mice^40^ were a gift from Drs. Cubillos Ruiz and Glimcher. C57BL/6-Tg(Tcra2D2, Tcrb2D2)1Kuch/J mice^72^ (The Jackson Laboratory, #006912) were used for co-coculture with DCs. All mice were housed in sterile autoclaved cages with irradiated food and acidified, autoclaved water. Mouse handling and weekly cage changes were performed by staff wearing sterile gowns, masks and gloves in a sterile biosafety hood. All mice were kept in a specific pathogen-free facility at the Hale Building of Transformative Medicine at the Brigham and Women’s Hospital. All mice were 8-10 weeks old at the time of EAE induction. All procedures were reviewed and approved under the IACUC guidelines at Brigham and Women’s Hospital.

### EAE

EAE was induced as described^71, 73^, using 150 µg of MOG_35-55_ (Genemed Synthesis Inc., #110582) mixed with complete Freund’s Adjuvant (20 mL Incomplete Freund’s Adjuvant (BD Biosciences, #BD263910) mixed with 100 mg *M. Tuberculosis* H-37Ra (BD Biosciences, #231141) at a ratio of 1:1 (v/v at a concentration of 5 mg/mL). All mice received 2 subcutaneous injections of 100 µL each of the MOG/CFA mix. All mice then received an intraperitoneal injection of *Pertussis* toxin (List Biological Laboratories, #180) at a concentration of 2 ng/µL in 200 µL of PBS at the day of EAE induction and 2 days later. Mice were monitored daily and scored as follows: 0 – no signs, 1 – fully limp tail, 2 – hindlimb weakness, 3 – hindlimb paralysis, 4 – forelimb paralysis, 5 – moribund. Mice were randomly assigned to treatment groups. All mice were scored blind to genotype.

### Isolation of mouse CNS cells

CNS cells were isolated by flow cytometry as described^71, 73^. In brief, mice were perfused with 1X PBS, the CNS was isolated, finely minced, digested in 0.66 mg/mL Papain (Sigma-Aldrich, #P4762) in HBSS for 20 min and then incubated for 20 min with and equal volume of DMEM with Collagenase D (Roche, #11088858001) and DNase I (Thermo Fisher Scientific, #90083) at 0.66 mg/mL and 8 U/mL respectively. Samples were shaken at 80rpm at 37°C. Tissue was mechanically dissociated using a 5 mL serological pipette and filtered through a 70 µm cell strainer (Fisher Scientific, #22363548) into a 50 mL conical. Tissue was centrifuged at 500g for 5 minutes and resuspended in 10 mL of 30% Percoll solution (9 mL Percoll (GE Healthcare Biosciences, #17-5445-01), 3 mL 10X PBS, 18 mL ddH2O). The Percoll suspension was centrifuged at 500g for 25 minutes with no brakes. Supernatant was discarded and the cell pellet was washed 1X PBS, centrifuged at 500g for 5 minutes and prepared for downstream applications.

### Isolation of mouse splenic cells

Spleens were incubated for 20 min with 2 mg/mL collagenase D (Roche, #11088858001) and then mashed through a 70-μm cell strainer. Red blood cells were lysed with ACK lysing buffer (Life Technology, #A10492-01) for 5 min, and then washed with 0.5% BSA, 2 mM EDTA pH=8.0 in 1X PBS.

### Salmonella Culturing and Infection

A modified *S. Typhimurium* LVS strain BRD509 expressing a short peptide sequence recognizable by 2SW1 tetramer staining (generously provided by Dr. Stephen McSorley) was used for all experiments. A stab of bacterial glycerol stock was first cultured overnight in 3mL Luria-Bertani (LB) broth at 37 °C in a shaking incubator at 200 rpm. The next day a subculture was prepared using 30 μl of the original culture in 3mL LB broth at 37 °C and 200 rpm for 4hrs or until the OD600 was in logarithmic range (0.6-0.9). CFU concentration was calculated using the conversion factor 1.0 (OD600) = 1×10^9^ CFU/ml. For infections, the bacteria culture was directly diluted to 1.25×10^7^ CFU/mL in sterile PBS and 200μL was injected via tail vein. Doses were validated by serial dilution and plating on LB agar plates at 37 °C then counting CFUs 12hrs later. Fourteen days post infection liver and cecum were weighted, then small pieces were weighed and used for CFU analysis. Tissue pieces were homogenized in 1mL sterile PBS using the gentleMACS Dissociator (Miltenyi Biotec), then homogenates were serially diluted and plated in triplicate on LB agar plates incubated overnight at 37°C. CFU counts were calculated per mg tissue, then extrapolated to total tissue burden using total tissue weights.

### Isolation of colon infiltrating cells

Briefly, tissue was cleaned, cut on 1cm pieces and incubated with 20 mL of pre-warmed PBS containing 1mM DTT (Thermo Fisher Scientific, #P2325), 2.5 mM EDTA and 2.5% FBS for 20 min at 37C. Then, samples were rinsed with PBS and 20 mL of PBS containing 5mM EDTA and 5% FBS was added for 20 min at 37C. Samples were rinsed 3 times with PBS on nylon strainer, minced with scissors and digested with 1 mg/mL Collagenase VIII (Sigma-Aldrich, #C2139-5G), 40U DNAse (ThermoFisher Scientific, #90083) in complete RPMI for 40 min at 37C. Finally, samples were subsequently filtered on 100 micron and 40 um cell strainers and stained as described above. CD45^+^ cells were isolated by CD45 magnetic beads isolation kit (130-052-301, Miltenyi Biotec).

### Isolation of liver immune cells

Hepatic immune cells were isolated as described^74^. Briefly, livers were obtained in PBS (Gibco) and passed through 100 mm nylon meshes, erythrocytes were removed using ACL lysis buffer. Hepatocytes were separated from immune cells by a 35% and 70% bilayer Percoll gradient (GE Healthcare) and used.

### T-cell FACS analysis

Cells were re-stimulated in RPMI containing 10% Fetal bovine serum (Gibco), 500 ng/mL ionomycin (Sigma-Aldrich), 500 ng/mL PMA (Sigma-Aldrich) GolgiPlug and BD GolgiStop (BD Bioscience), incubated at 37 C and 5% CO_2_ for 4h. Re-stimulated cells were then washed with PBS, and incubated for 30 min with Dasatanib (5 nM, Sigma-Aldrich), and purified Fc block (553142, BD Biosciences) followed by a 1 h incubation with APC-labeled S2W1 tetramer (TS-M722-2, MBL International). Tetramer-stained cells were washed and stained using the following antibodies: BUV661 labeled anti-CD45 antibody (612975, BD Biosciences), BV650 labeled anti-CD3 antibody (740530, BD Biosciences), PE-Cy7 labeled anti-CD4 antibody (100422, 100547, Biolegend). Cells were then fixed and permeabilized using the Foxp3 Fixation/Permeabilization kit (eBioscience). Intracellular antibodies were: APC/Cy7 anti-mouse IFN-γ (561479, BD Biosciences); PE anti-mouse IL-17A (506904, BioLegend). Cells were acquired on a Symphony A5 (BD Biosciences) and analyzed on Flowjo 10 (Becton Dickison).

### Bone marrow-derived dendritic cell (BMDC) differentiation

To obtain bone marrow cells, femurs and tibiae from C57BL/6J mice were removed and obtained by flush with 1 mL of with 0.5% BSA, 2 mM EDTA pH=8.0 in 1X PBS. Red blood cells were lysed with ACK lysing buffer (Life Technology, #A10492-01) for 5 min and washed with 0.5% BSA, 2 mM EDTA pH=8.0 in 1X PBS. For dendritic cells differentiation, cells were cultured in non-adherent petri dishes (100 mm) at 1×10^6^ cells/mL in DMEM/F12+GlutaMAX (Life Technologies, #10565042) supplemented with 10% FBS (Life Technologies, #10438026), 1% penicillin/streptomycin (Life Technologies, #15140122), 1% Sodium Pyruvate (Life Technologies, #11360070), 1% Hepes (Life Technologies, #15630106), and 1% MEM non-essential amino acids (Life Technologies, #11140050) containing 20 ng/mL of GM-CSF (PeproTech, #315-03). The media was replaced at day 3 and 5, and cells were completely differentiated at day 8 and prepared for downstream applications.

### DC stimulation

DCs were treated for 6h with 100 ng/mL Ultrapure LPS, E.coli 0111:B4 (tlrl-3pelps, InvivoGen,), 0.1, 1 or 10 mM Sodium L-Lactate (71718, Sigma Aldrich,), 1 or 10 mM Sodium D-Lactate (71716, Millipore Sigma,), 1.5 mM Pyruvate Solution (103578-100, Agilent Technologies,), 20 μM MitoParaquant (MitoPQ) (ab146819, Abcam,), 50 μM Mito-TEMPO (SML0737, Sigma Aldrich), 0.1, 0.2 or 1 mM HIF-1α inhibitor (5655, Tocris Bioscience), 6.5 μM Dimethyl Fumarate (DMF) (242926, Sigma Aldrich), 10 mM Dimethylmalonate (136441-250G, Sigma-Aldrich), 5 mM Diethyl Succinate (112402, Sigma Aldrich), 125μM 4-Octyl Itaconate (3133-16-2, Cayman Chemical Company), 1mM Dimethyl α-ketoglutarate (349631-5G, Sigma Aldrich), 10μM ML228 (4565, Tocris Bioscience), 500 μM deferoxamine (D9533-1G, Sigma Aldrich). In some studies BMDCs were pre-treated 1h before stimulation with: 2mM Sodium Oxamate (02751, Sigma Aldrich), 3 mM sodium 3-hydroxibutyrate as GPR81 antagonist (54965-10G-F, Sigma-Aldrich) and 0.5 μM AR-C 141990 hydrochloride (5658, Tocris) or 100 nM AZD3965 (HY-12750, MedChem Express) as MCT-1 inhibitors. For cultures under hypoxia, cells were incubated for 6h in hypoxia incubator chamber (Stem cell Technologies, #27310) at 37°C and 1% CO_2_.

### Human DC and T cell cultures

PBMCs were isolated from healthy blood donors by Ficoll-Paque PLUS density gradient (Sigma Aldrich, GE17-1440-03). The protocols for this study received prior approval from all institutional review boards, and informed consent was obtained from each subject. Exclusion criteria included pregnancy and antibiotics or vaccines within the past 3 months. DCs were enriched using Pan-DC enrichment kit (130-100-777, Miltenyi), enriched DCs were stained with BV421 anti-human CD11c (301628, Biolegend), APC Cy7 anti-human HLA-DR (307618, Biolegend) and FITC anti-human CD3 (555232, BD Biosciences). T cells were sorted as CD3^+^ cells. DCs sorted as CD3^neg^CD11c^+^HLA-DR^+^ were silenced and 48h cells were stimulated with LPS (100ng/mL) and 10 µg/mL of OVA 323-339 (SP-51023-1, Genemed) in the presence of D-LA or L-LA (1mM) or vehicle. After 6h, cells were washed, lysed to evaluate cytokine gene expression or co-cultured with allogenic T cells in a 1:10 ratio for 48h. HIF1a was silenced on DCs using Accell Human *HIF1a*-siRNA smart pool (E-004018-00-0005, Dharmacon,). Then, DC:T cell co-cultures were performed in the presence of soluble anti-human CD3 (16-0037-85, ThermoFisher).

### Knockdown with siRNA

siRNA pool was mixed with Interferin (Polyplus-transfection, #409-10) in Opti-MEM (Life Technologies, #31985062), incubated for 10min at room temperature and added to pre-plated 200,000 BMDCs. 48h later, cells were used for downstream assays. siRNA pools (1 nM) used were non-targeting siRNA (Dharmacon, #D-001810-10-05), si*Hif1a* (L-040638-00-0005, Dharmacon), si*Vhl* (E-040755-00-0005,Dharmacon), si*Slc16a1* (L-058863-01-0005, Dharmacon), and si*Ndufa4l2* (L-160257-00-0005, Dharmacon). Knockdown efficiency was confirmed by qPCR.

### DC-T-cell co-culture

3×10^4^ DCs were plated in Costar low attached 96-well plate (Sigma-Aldrich, #CLS3474-24EA) and stimulated with LPS or LPS + L-LA or D-LA in presence of 20 μg/mL MOG_35-55_ for 6h. Next, cells were washed with DMEM/F12+GlutaMAX and co-cultured with 3×10^5^ naïve 2D2 CD4^+^ T-cells for 48h.

### Recall response

For recall response, 4×10^5^ splenocytes per well were cultured in complete RPMI medium for 72h in 96-well plates in the presence of 5, 20 or 100 μg/mL of MOG_35–55_ (Genemed Synthesis, #110582). During the last 16h, cells were pulsed with 1 μCi [3H]-thymidine (PerkinElmer, #NET027A005MC) followed by collection on glass fiber filters (PerkinElmer, #1450-421) and analysis of incorporated [3H]-thymidine in a β-counter (PerkinElmer, #1450 MicroBeta TriLux).

### ELISA

Supernatants from splenocytes stimulated with 20 μg/mL of MOG_35–55_ peptide (Genemed Synthesis, #110582) for 72 hours, or from BMDCs-CD4^+^ T cells co-cultures were collected to quantify cytokine release by ELISA following manufacturer’s instruction. Briefly, Costar 96-well plates (Corning, #3690) were coated with capture antibodies diluted in 1X PBS: anti-mouse GM-CSF Capture (Invitrogen, #88-7334-88, 1:250), anti-mouse IL-10 capture (Invitrogen, #88-7105-88, 1:250), purified anti-mouse IFN-γ (BD Biosciences, #551309, 1:500), and purified anti-mouse IL-17A (BD Biosciences, #555068, 1:500) overnight at 4C. Plates were washed 2X with 0.05% Tween in 1X PBS (Boston BioProducts, #IBB-171X) and blocked with 1% BSA in 1X PBS (Thermo Fisher Scientific, #37525) for 2 hours at room temperature. Standard curve was prepared from cytokine standards diluted in 1X Elispot diluent (eBioscience, #00-4202-56). Samples were diluted 4X for IFN-γ, 2X for GM-CSF and IL-17A in Elispot diluent and plated pure for IL-10. Plates were washed 2X with 0.05% Tween in 1X PBS and samples and standard curve were added and incubated overnight at 4C. The next day, plates were washed 5X with 0.05% Tween in 1X PBS and incubated with detection antibodies diluted in 1X Elispot diluent: anti-mouse GM-CSF Detection (Invitrogen, #88-7334-88, 1:250), anti-mouse IL-10 detection (Invitrogen, #88-7105-88, 1:250), biotin anti-mouse IFN-γ (BD Biosciences, #554410, 1:500), and biotin anti-mouse IL-17A (BD Biosciences, #555067, 1:500) shaken for 1 h at room temperature. Afterwards, plates were washed 8X with 0.05% Tween in 1X PBS and incubated with Avidin-HRP diluted in 1X Elispot diluent (Invitrogen, #88-7334-88, 1:250) shaken for 1 h at room temperature. Next, plates were washed 8X with 0.05% Tween in 1X PBS and revealed using 1X TMB Substrate Solution (Invitrogen, #00-4201-56). The reaction was stopped by KPL TBM Stop Solution (SeraCare, #5150-0021) and plates were read at 450nm on GloMax Explorer Multimode Microplate Reader (Promega).

### Mitochondrial ROS production

5×10^5^ BMDCs were stimulated with LPS, L-LA, LPS + L-LA and vehicle for 1h at 37C. Then, cells were stained with 5 μM of mitoSOX mitochondrial indicator (Thermo Fisher Scientific, #M36008) for 10 min at 37°C. Next, cells were washed 3X with prewarmed HBSS and fluorescence was read on GloMax Explorer Multimode Microplate Reader (Promega).

### Flow cytometry of DCs

To analyze DCs by flow cytometry, CNS, lymph nodes and splenic cell suspensions were incubated with surface antibodies and a live/dead cell marker on ice. After 30 min, cells were washed with 0.5% BSA, 2mM EDTA in 1X PBS and fixed according to the manufacturer’s protocol (eBiosciences, #00-5523-00). Intracellular staining was performed for 1h at room temperature. Surface antibodies used in this study were: BUV395 anti-mouse MHC-II (17-5321-82, Invitrogen, 1:200); BUV496 anti-mouse CD24 (564664, BD Biosciences, 1:100); BUV563 anti-mouse Ly-6G (612921, BD Biosciences, 1:100); BUV661 anti-mouse CD45 (103147, BioLegend, 1:100); BV570 anti-mouse Ly-6C (128030, BioLegend, 1:100); BV605 anti-mouse CD80 (563052, BD Biosciences, 1:100); BV650 anti-mouse CD3 (100229, BioLegend, 1:100); BV711 anti-mouse CD8a (100748, BioLegend, 1:100); BV786 anti-mouse CD11b (101243, BioLegend, 1:100); PE-Texas Red anti-mouse CD11c (117348, BioLegend, 1:100); PeCy7 anti-mouse CD4 (100422, BioLegend, 1:100); APC anti-mouse/human CD45R/B220 (103212, BioLegend, 1:100); APC-R700 anti-mouse CD103 (565529, BD Biosciences, 1:100); APC/Cy7 anti-mouse F4/80 (123118, BioLegend, 1:100), Alexa 647 labeled anti-mouse CD36 (102610, Biolegend, 1:200). Intracellular antibody used was Alexa Fluor 488 anti-mouse HIF-1α (BS-0737R-A488, Bioss Antibodies, 1:100). FACs was performed on a Symphony A5 (BD Biosciences).

### FACS analysis of T-cells

To analyze T-cells, CNS, lymph nodes and splenic cell suspensions were stimulated with 50 ng/mL phorbol 12-myristate 13-acetate (PMA, Sigma-Aldrich, #P8139), 1 µM Ionomycin (Sigma-Aldrich, #I3909-1ML), GolgiStop (BD Biosciences, #554724, 1:1500) and GolgiPlug (BD Biosciences, #555029, 1:1500) diluted in RPMI (Life Technologies, #11875119) containing 10% FBS, 1% penicillin/streptomycin, 50 µM 2-metcaptoethanol (Sigma-Aldrich, #M6250), and 1% non-essential amino acids (Life Technologies, #11140050). After 4 hours, cell suspensions were washed with 0.5% BSA, 2 mM EDTA in 1X PBS and incubated with surface antibodies and a live/dead cell marker on ice. After 30 min, cells were washed with 0.5% BSA, 2mM EDTA in 1X PBS and fixed according to the manufacturer’s protocol of an intracellular labeling kit (00-5523-00, eBiosciences). Surface antibodies used in this study were: BUV661 anti-mouse CD45 (103147, BioLegend, 1:100); PeCy7 anti-mouse CD4 (100422, BioLegend, 1:100); BV750 anti-mouse CD3 (100249, BioLegend, 1:100); BUV395 anti-mouse CD69 (740220, BD Biosciences, 1:100); BUV737 anti-mouse CD11b (564443, BD Biosciences, 1:100); BV805 anti-mouse CD8a (612898, BD Biosciences, 1:100); PE/Cy5 anti-mouse CD44 (103010, BioLegend, 1:100). Intracellular antibodies were: BV421 anti-mouse GM-CSF (564747, BD Biosciences, 1:100); APC/Cy7 anti-mouse IFN-γ (561479, BD Biosciences, 1:100); PE anti-mouse IL-17A (506904, BioLegend, 1:100). Cells were acquired on a Symphony A5 (BD Biosciences) and analyzed on Flowjo 10 (Becton Dickison).

### RNA isolation

Sorted DCs were lysed in extraction buffer and RNA was isolated following manufacturer’s instruction (Thermo Fisher Scientific, # KIT0204). BMDCs were lysed in RLT Buffer and RNA was isolated using the Qiagen RNeasy Mini kit (Qiagen, #74106). cDNA was transcribed using the High-Capacity cDNA Reverse Transcription Kit (Life Technologies, #4368813). Gene expression was measured by qPCR using Taqman Fast Universal PCR Master Mix (Life Technologies, #4367846). Taqman probes used in this study are: *Gapdh* (Mm99999915_g1), *Il1b* (Mm00434228_m1), *Il6* (Mm00446190_m1), *Il12* (Mm00434169_m1), *Il23* (Mm00518984_m1), *Tnf* (Mm00443528_m1), *Ndufa4l2* (Mm01160374_g1), *sXbp1* (Mm03464496_m1), *Xbp1* (Mm00457357_m1), *Gclm* (Mm01324400_m1), *Gls* (Mm01257297_m1), *Phgdh* (Mm01623589_g1), *Shmt2* (Mm00659512_g1), *Slc7a11* (Mm00442530_m1), *Lonp1* (Mm01236887_m1), *Cox4i1* (Mm0250094_m1), *Cox4i2* (Mm00446387_m1), *Hif1a* (Mm00468869_m1), *Cd36* (Mm00432403_m1), *Vhl* (Mm00494137_m1), *Mct1* (Mm01306379_m1), *Slc2a1* (Mm00441480_m1), *GAPDH* (Hs02786624_g1), *IL23* (Hs00372324_m1), *IL1B* (Hs01555410_m1), *IL6* (Hs00174131_m1), *HIF1a* (Hs00153153_m1), *IFNG* (Hs00989291_m1), *CSF2* (Hs00929873_m1). qPCR data were analyzed by the ddCt method by normalizing the expression of each gene to *Gapdh* and then to the control group.

### Fluorescent Glucose Uptake

2×10^4^ DCs were stimulated with LPS (100ng/mL) in glucose free RPMI (11-879-020, Fisher Scientific). After 1 h, cells were washed and stained with 10 μΜ 2NBDG (N13195, Thermofisher) at 37C. After 20 min, samples were acquired in a BD LSR Fortessa.

### Pyruvate quantification

Pyruvate levels were quantified in BMDC lysates after 1h stimulation using EnzyChrom pyruvate assay kit (EPYR-100, BioAssay Systems), followed the procedure suggested by the manufacturer. The signal was detected on GloMax microplate reader (Promega).

### LA incorporation

2×10^6^ BMDCs were treated with 1mM uniformly labeled 13C lactate (L-LA*) (CLM-1579-N-0.1MG, Cambridge Isotope Laboratories), in the presence or absence of LPS (100ng/mL). After 1h, cells were washed with cold 1X and then lysed with 80% of HPLC grade methanol (34860, Sigma-Aldrich). Analysis of 13C incorporation into TCA intermediates was performed at the Broad Institute.

### Chromatin immunoprecipitation (ChIP)

Approximately 2 million BMDCs were processed according to the ChIP-IT Express Enzymatic Shearing and ChIP protocol (Active Motif, #53009) as previously described^75^. Briefly, cells were fixed in 1% formaldehyde for 10 minutes with gentle agitation, washed in 1X PBS, washed for 5 minutes in 1X glycine stop-fix solution in PBS, and scraped in 1X PBS supplemented with 500 µM PMSF. Cells were pelleted, nuclei isolated, and chromatin sheared using the Enzymatic Shearing Cocktail for 10 minutes at 37C. Sheared chromatin was immunoprecipitated according to the Active Motif protocol overnight at 4°C, with rotation. The next day, the protein-bound magnetic beads were washed with ChIP buffer 1, ChIP buffer 2, and finally with 1X TE. Cross-links were reversed in 100 µL of 0.1% SDS and 300 mM NaCl in 1X TE at 65C for 4-5 hours. DNA was purified using QIAquick PCR Purification Kit (Qiagen, #28104). qPCR was performed using Fast SYBR Green Master Mix (Thermo Fisher Scientific, #4385612). Anti-IgG immunoprecipitation and input were used as controls. Antibodies used were mouse Xbp-1 (Santa Cruz Biotechnology, #SC-8015) and IgG1 isotype control (Abcam, #ab171870). PCR primers were designed with Primer3 to generate 50-150 bp amplicons. Primer sequences used: XBP1-IL1B site 1 F: 5’-CCTGGGCTCTCTGAGTTAGCAGTCTAGTGA-3’; R: 5’- AGTGCAAAGGTGGTGAACGAGGCATCTG-3’; XBP1-IL1B site 2 F: 5’- GGCTTGCTTCCAGAGTTCCCTGACCCTAT-3’; R: 5’- GAGTTTGGTAACTGGATGGTGCTTCTGTGC-3’; XBP1-IL6 site 1 F: 5’- CCTGCGTTTAAATAACATCAGCTTTAGCTT-3’; R: 5’- GCACAATGTGACGTCGTTTAGCATCGAA-3; XBP1-IL23 site 1 F: 5’- GCTTCCAACCCTCCAGATCC-3’; R: 3’-ACCTTCCCAGTCCTCCAAGT-5’; XBP1-IL23 site 2 F: 5’-CCTCTAGCCACAACAACCTC-3’; R: 3’-CCTTCACACTAGCAGGTGACT- 5’. Data were analyzed by ddCt relative to IgG control or input DNA.

### Oxygen consumption rate

3.5×10^5^ *Hif1a*-silenced, *Ndufa4l2*-silenced, non-targeted BMDCs or HIF-1α-deficient BMDCs were plated in Seahorse XF96 Cell Culture Microplates (Agilent Technologies, #101085-004) and stimulated with 10 mM of L-LA, 1.5 mM of Pyruvate, or vehicle for 6h. Oxygen consumption rate (OCR) was evaluated after 2 µM Oligomycin, 1.5 µM FCCP and 1 µM Rotenone/1 µM Antimycin A from Seahorse XF Cell Mito Stress Test (103015-100, Agilent Technologies), and read using Seahorse XFe96 Analyzer (Agilent Technologies). OCR were normalized to total cell number evaluated with CyQUANT cell proliferation assay kit (C7026, Invitrogen).

### Extracellular acidification rate

3.5×10^5^ HIF-1α-deficient or WT BMDCs were plated in Seahorse XF96 Cell Culture Microplates (Agilent Technologies, #101085-004). Cells were kept in 180 μL of assay medium (supplemented with 2 mM glutamine, pH 7.4) 1h. ECAR was determined at the beginning of the assay and after the sequential addition of 10 mM d-glucose, 1 μM oligomycin, and 100 mM 2-DG. ECAR were normalized to total cell number evaluated with CyQUANT cell proliferation assay kit (C7026, Invitrogen).

### Luciferase assay

4×10^6^ BMDCs from FVB.129S6-Gt(ROSA)26Sortm2(HIF1A/luc)Kael/J mice mice (006206, The Jackson Laboratory,) were stimulated with LPS or LPS + L-LA (10mM) for 6h. Next, the luciferase activity was quantify using Dual-Luciferase Reporter Assay System (Promega, #E1910) according to the manufacturer’s protocol. In another set of experiments, HepG2 (ATCC HB-8065) or DC2.4 (SCC142, Millipore) cells were cultured in DMEM supplied with 10% FBS (GIBCO) and Penicillin/Streptomycin (Thermo Fisher Scientific). Ppar alpha, Ndufa4l2, Hif1a promoter reporter plasmids expressing Gaussia luciferase were acquired from GeneCopoeia. To evaluate Ndufa4l2 promoter activity cells were stimulated with vehicle, L-LA (10mM), LPS (100ng/ml) or the combination of L-LA and LPS. To evaluate Ppar alpha and Hif1a promoter activity, cells were stimulated with Fenofibrate, L-LA (10mM) or D-LA (10mM). For over-expression assays, HIF1a plasmid (OHu27176, GenScript), Ndufa4l2 plasmid (EX-J0135-M02, GeneCopoeia) or Empty control vector (GeneCopoeia) were used. All plasmids were transfected using Lipofectamine 2000 (11668019, Thermo Fisher Scientific) following manufacturer’s instructions. Luciferase activity was evaluated with Gaussia Luciferase Flash Assay Kit (16159, Thermo Fisher Scientific).

### Lactate administration

L-LA or D-LA (125 mg/kg, daily) was administered ip, nasal, or iv starting 3 days before EAE induction. Tissues were harvested 28 days after EAE induction and cell suspensions were used for downstream applications.

### Engineering a D-lactate producing live therapeutic

Strains used in this work were listed in **Table 2**. Escherichia coli Nissle 1917 (EcN), designated as SYN001 here, was originally purchased from the German Collection of Microorganisms and Cell Cultures (DSMZ Braunschweig, E. coli DSM 6601). EcN (SYN001) with strep resistance was generated by streaking ∼ e^11^ cells of SYN001 on LB plate containing 300 ng/mL streptomycin followed by taking the single colony formed to have mutated to be strep resistant and designated as EcN-StrpR (SYN094). Pta gene on the pyruvate-acetate pathway was deleted to limit the carbon flux through acetate producing using the lambda red recombineering technique as described^69^ resulting in strain EcN-Δpta. The gene encoding E. coli LdhA protein under the control of Ptemp promoter was subcloned into a Synlogic vector containing ampicillin resistant gene, a low copy number origin of replication pSC101. The resulting plasmid was used to electroporate EcN-Δpta resulting in strain EcN-Lactate.

### Engineering a reporter strain using green fluorescent protein

The gene encoding a green fluorescent protein (GFP) together with an upstream constitutive promoter J23100 was synthesized and subcloned into a Synlogic vector containing ampicillin resistant gene, a low copy number origin of replication pSC101, resulting in plasmid pSC101-Pconst-rfp-ampR. Then the gene encoding a green fluorescent protein under the control of the heat inducible promoter cassette was synthesized and subcloned into this plasmid pSC101-Pconst-rfp-ampR to obtain the heat inducible GFP reporter plasmid pSC101-Ptemp-gfp-Pconst-rfp-ampR. This final plasmid is used to transformed into EcN to produce EcN-GFP.

### Growth and induction of strains in bioreactors

Cells were inoculated in 50 mL of FM1 medium supplemented with 25 g/L Glucose in a 500-mL Ultra-Yield flask (Thomson). Cells were grown at 37 °C with shaking at 350 rpm overnight. Next day, 110 mL of the overnight culture was used to inoculate 4 L of FM1 in an Eppendorf BioFlow 115 bioreactor (starting OD600 of ∼0.5). The fermenter was controlled at 60% dissolved oxygen and 30 °C with agitation, air, and oxygen supplementation, and controlled to pH 7 using ammonium hydroxide. The expression of ldhA gene was induced at OD600 ∼ 5 by changing the reactor temperature from 30 °C to 37 °C for 2 hours. Cells were harvested after 2 hours and at OD600 ∼20. Cells were harvested by centrifugation at 4,500 g for 30 min at 4°C, resuspended in formulation buffer, and stored at −80 °C. OD600 was measured on a BioPhotometer Plus spectrophotometer (Eppendorf).

### Protein seq of ldhA enzyme used in plasmid to build EcN-Lactate strain

MKLAVYSTKQYDKKYLQQVNESFGFELEFFDFLLTEKTAKTANGCEAVCIFVNDDGSRPVLEELKKHGVKYIALRCAGFNNVDLDAAKELGLKVVRVPAYDPEAVAEHAI GMMMTLNRRIHRAYQRTRDANFSLEGLTGFTMYGKTAGVIGTGKIGVAMLRILKGFGMRLLAFDPYPSAAALELGVEYVDLPTLFSESDVISLHCPLTPENYHLLNEAAFEQMKNGVMIVNTSRGALIDSQAAIEALKNQKIGSLGMDVYENERDLFFEDKSNDVIQDDVFRRLSACHNVLFTGHQAFLTAEALTSISQTTLQNLSNLEKGETCPNELV

### DNA sequence of plasmid pSC101-Ptemp-ldhA-ampR

aaaaatgaagttttaaatcaatctaaagtatatatgagtaaacttggtctgacagttaccaatgcttaatcagtgaggcacctatctcagcgatctgtctatttcgttcatccatagttgcctgactccccgtcgtgtagataactacgatacgggagggcttaccatctggccccagtgctgcaatgataccgcgagaaccacgctcaccggctccagatttatcagcaataaaccagccagccggaagggccgagcgcagaagtggtcctgcaactttatccgcctccatccagtctattaattgttgccgggaagctagagtaagtagttcgccagttaatagtttgcgcaacgttgttgccattgctacaggcatcgtggtgtcacgctcgtcgtttggtatggcttcattcagctccggttcccaacgatcaaggcgagttacatgatcccccatgttgtgcaaaaaagcggttagctccttcggtcctccgatcgttgtcagaagtaagttggccgcagtgttatcactcatggttatggcagcactgcataattctcttactgtcatgccatccgtaagatgcttttctgtgactggtgagtactcaaccaagtcattctgagaatagtgtatgcggcgaccgagttgctcttgcccggcgtcaatacgggataataccgcgccacatagcagaactttaaaagtgctcatcattggaaaacgttcttcggggcgaaaactctcaaggatcttaccgctgttgagatccagttcgatgtaacccactcgtgcacccaactgatcttcagcatcttttactttcaccagcgtttctgggtgagcaaaaacaggaaggcaaaatgccgcaaaaaagggaataagggcgacacggaaatgttgaatactcatactcttcctttttcaatattattgaagcatttatcagggttattgtctcatgagcggatacatatttgaatgtatttagaaaaataaacaaataggggttccgcgcacatttccccgaaaagtgccacctgacgtctaagaaaccattattatcatgacattaacctataaaaataggcgtatcacgaggccctttcgtctcgcgcgtttcggtgatgacggtgaaaacctctgacacatgcagctcccggagacggtcacagcttgtctgtaagcggatgccgggagcagacaagcccgtcagggcgcgtcagcgggtgttggcgggtgtcggggctggcttaactatgcggcatcagagcagattgtactgagagtgcaccatatgcggtgtgaaataccgcacagatgcgtaaggagaaaataccgcatcaggcgccattcgccattcaggctgcgcaactgttgggaagggcgatcggtgcgggcctcttcgctattacgccagctggcgaaagggggatgtgctgcaaggcgattaagttgggtaacgccagggttttcccagtcacgacgttgtaaaacgacggccagtgcgCTCCCGGAGACGGTCACAGCTTGTCaaaaaaaaaccccgcttcggcggggtttttttttGGTACCTCATCAGCCAAACGTCTCTTCAGGCCACTGACTAGCGATAACTTT CCCCACAACGGAACAACTCTCATTGCATGGGATCATTGGGTACTGTGGGTTTAGTGGTTGTAAAAACACCTGACCGCTATCCCTGATCAGTTTCTTGAAGGTAAACTCATCACCCCCAAGTCTGGCTATGCAGAAATCACCTGGCTCAACAGCCTGCTCAGGGTCAACGAGAATTAACATTCCGTCAGGAAAGCTTGGCTTGGAGCCTGTTGGTGCGGTCATGGAATTACCTTCAACCTCAAGCCAGAATGCAGAATCACTGGCTTTTTTGGTTGTGCTTACCCATCTCTCCGCATCACCTTTGGTAAAGGTTCTAAGCTTAGGTGAGAACATCCCTGCCTGAACATGAGAAAAAACAGGGTACTCATACTCACTTCTAAGTGACGGCTGCATACTAACCGCTTCATACATCTCGTAGATTTCTCTGGCGATTGAAGGGCTAAATTCTTCAACGCTAACTTTGAGAATTTTTGTAAGCAATGCGGCGTTATAAGCATTTAATGCATTGATGCCATTAAATAAAGCACCAACGCCTGACTGCCCCATCCCCATCTTGTCTGCGACAGATTCCTGGGATAAGCCAAGTTCATTTTTCTTTTTTTCATAAATTGCTTTAAGGCGACGTGCGTCCTCAAGCTGCTCTTGTGTTAATGGTTTCTTTTTTGTGCTCATACGTTAAATCTATCACCGCAAGGGATAAATATCTAACACCGTGCGTGTTGACTATTTTACCTCTGGCGGTGATAATGGTTGCATaagtgaggatccaaagtgaactctagaaataattttgtttaactttaagaaggaggtatacatATGAAACTTGCTGTATATAGTACCAAACAGTACGACAAAAAGTACCTTCAACAGGTCAACGAGAGCTTTGGTTTCGAACTTGAATTTTTCGACTTTTTACTTACCGAGAAAACGGCAAAAACGGCGAACGGATGTGAAGCGGTTTGCATTTTCGTCAACGACGACGGCAGCCGCCCTGTTTTAGAAGAGTTAAAGAAACATGGAGTTAAATACATCGCATTACGTTGTGCAGGTTTCAACAACGTTGATCTGGATGCTGCGAAGGAACTGGGATTGAAAGTTGTGCGCGTGCCCGCTTATGACCCAGAGGCGGTTGCGGAACACGCTATTGGTATGATGATGACCCTTAATCGTCGCATCCATCGTGCATATCAGCGCACGCGCGATGCTAACTTCAGTTTAGAAGGATTAACGGGATTTACAATGTACGGGAAGACCGCTGGCGTGATTGGCACCGGAAAAATCGGTGTGGCAATGCTGCGTATCTTGAAGGGGT TTGGCATGCGTTTGTTAGCATTTGATCCCTATCCAAGTGCCGCGGCCCTGGAACTGGGAGTGGAATATGTTGATTTGCCAACTTTGTTTAGCGAGTCCGATGTTATCTCATTGCATTGTCCACTTACTCCGGAGAATTATCATTTATTGAATGAAGCCGCCTTCGAACAAATGAAAAATGGAGTGATGATCGTAAATACAAGTCGTGGCGCGTTGATCGATTCGCAGGCAGCGATCGAAGCGTTAAAAAATCAAAAGATTGGATCACTGGGCATGGATGTCTATGAAAACGAGCGCGACCTTTTCTTTGAAGACAAAAGTAATGATGTTATCCAAGATGATGTATTTCGCCGTCTGTCGGCATGCCATAATGTACTTTTTACGGGTCACCAAGCATTCCTTACTGCCGAGGCTCTGACTAGCATTTCACAAACCACTCTTCAGAATCTTTCAAATCTTGAGAAAGGTGAGACGTGCCCCAATGAATTGGTTtaaGCATGCTAATCAGCCGTGGAATTCGGTCTCaGGAGgtacgcatggcatggatgaccgatggtagtgtgggctctccccatgcgagagtagggaactgccaggcatcaaataaaacgaaaggctcagtcgaaagactgggcctttcgttttatctgttgtttgtcggtgaacgctctcctgagtaggacaaatccgccgggagcggatttgaacgttgcgaagcaacggcccggagggtggcgggcaggacgcccgccataaactgccaggcatcaaattaagcagaaggccatcctgacggatggcctttttgcgtggccagtgccaagcttgcatgcgtgccagctgcattaatgaagaaatcatgctggaagaataacagctcactcaaaggcggtagtacgggttttgctgcccgcaaacgggctgttctggtgttgctagtttgttatcagaatcgcagatccggcttcagccggtttgccggctgaaagcgctatttcttccagaattgccatgattttttccccacgggaggcgtcactggctcccgtgttgtcggcagctttgattcgataagcagcatcgcctgtttcaggctgtctatgtgtgactgttgagctgtaacaagttgtctcaggtgttcaatttcatgttctagttgctttgttttactggtttcacctgttctattaggtgttacatgctgttcatctgttacattgtcgatctgttcatggtgaacagctttgaatgcaccaaaaactcgtaaaagctctgatgtatctatcttttttacaccgttttcatctgtgcatatggacagttttccctttgatatgtaacggtgaacagttgttctacttttgtttgttagtcttgatgcttcactgatagatacaagagccataagaacctcagatccttccgtatttagccagtatgttctctagtgtggttcgttgtttttgcgtgagccatgagaacgaaccattgagatcatacttactttgcatgtcactcaaaaattttgcctcaaaactggtgagctgaatttttgcagttaaagcatcgtgtagtgtttttcttagtccgttatgtaggtaggaatctgatgtaatggttgttggtattttgtcaccattcatttttatctggttgttctcaagttcggttacgagatccatttgtctatctagttcaacttggaaaatcaacgtatcagtcgggcggcctcgcttatcaaccaccaatttcatattgctgtaagtgtttaaatctttacttattggtttcaaaacccattggttaagcctttta aactcatggtagttattttcaagcattaacatgaacttaaattcatcaaggctaatctctatatttgccttgtgagttttcttttgtgttagttcttttaataaccactcataaatcctcatagagtatttgttttcaaaagacttaacatgttccagattatattttatgaatttttttaactggaaaagataaggcaatatctcttcactaaaaactaattctaatttttcgcttgagaacttggcatagtttgtccactggaaaatctcaaagcctttaaccaaaggattcctgatttccacagttctcgtcatcagctctctggttgctttagctaatacaccataagcattttccctactgatgttcatcatctgagcgtattggttataagtgaacgataccgtccgttctttccttgtagggttttcaatcgtggggttgagtagtgccacacagcataaaattagcttggtttcatgctccgttaagtcatagcgactaatcgctagttcatttgctttgaaaacaactaattcagacatacatctcaattggtctaggtgattttaatcactataccaattgagatgggctagtcaatgataattactagtccttttcctttgagttgtgggtatctgtaaattctgctagacctttgctggaaaacttgtaaattctgctagaccctctgtaaattccgctagacctttgtgtgttttttttgtttatattcaagtggttataatttatagaataaagaaagaataaaaaaagataaaaagaatagatcccagccctgtgtataactcactactttagtcagttccgcagtattacaaaaggatgtcgcaaacgctgtttgctcctctacaaaacagaccttaaaaccctaaaggcttaagtagcaccctcgcaagctcgggcaaatcgctgaatattccttttgtctccgaccatcaggcacctgagtcgctgtctttttcgtgacattcagttcgctgcgctcacggctctggcagtgaatgggggtaaatggcactacaggcgccttttatggattcatgcaaggaaactacccataatacaagaaaagcccgtcacgggcttctcagggcgttttatggcgggtctgctatgtggtgctatctgactttttgctgttcagcagttcctgccctctgattttccagtctgaccacttcggattatcccgtgacaggtcattcagactggctaatgcacccagtaaggcagcggtatcatcaacaggcttacccgtcttactgtcttttctacggggtctgacgctcagtggaacgaaaactcacgttaagggattttggtcatgagattatcaaaaaggatcttcacctagatccttttaaatt

### DNA sequence of plasmid pSC101-Ptemp-gfp-Pconst-rfp-ampR

cAACTTAGGCGTAAACCTCGTTGCCACCTACTGTCAGACCAAGTTTACTCATATATACTTTAGATTGATTTAAAACTTCATTTTTAATTTAAAAGGATCTAGGTGAAGATCCTTTTTGATAATCTCATGACCAAAATCCCTTAACGTGAGTTTTCGTTCCACTGAGCGTCAGACCCCGTAGAAAAGACAGTAAGACGGGTAAGCCTGTTGATGATACCGCTGCCTTACTGGGTGCATTAGCCAGTCTGAATGACCTGTCACGGGATAATCCGAAGTGGTCAGACTGGAAAATCAGAGGGCAGGAACTGCTGAACAGCAAAAAGTCAGATAGCACCACATAGCAGACCCGCCATAAAACGCCCTGAGAAGCCCGTG ACGGGCTTTTCTTGTATTATGGGTAGTTTCCTTGCATGAATCCATAAAAGGCGCCTGTAGTGCCATTTACCCCCATTCACTGCCAGAGCCGTGAGCGCAGCGAACTGAATGTCACGAAAAAGACAGCGACTCAGGTGCCTGATGGTCGGAGACAAAAGGAATATTCAGCGATTTGCCCGAGCTTGCGAGGGTGCTACTTAAGCCTTTAGGGTTTTAAGGTCTGTTTTGTAGAGGAGCAAACAGCGTTTGCGACATCCTTTTGTAATACTGCGGAACTGACTAAAGTAGTGAGTTATACACAGGGCTGGGATCTATTCTTTTTATCTTTTTTTATTCTTTCTTTATTCTATAAATTATAACCACTTGAATATAAACAAAAAAAACACACAAAGGTCTAGCGGAATTTACAGAGGGTCTAGCAGAATTTACAAGTTTTCCAGCAAAGGTCTAGCAGAATTTACAGATACCCACAACTCAAAGGAAAAGGACTAGTAATTATCATTGACTAGCCCATCTCAATTGGTATAGTGATTAAAATCACCTAGACCAATTGAGATGTATGTCTGAATTAGTTGTTTTCAAAGCAAATGAACTAGCGATTAGTCGCTATGACTTAACGGAGCATGAAACCAAGCTAATTTTATGCTGTGTGGCACTACTCAACCCCACGATTGAAAACCCTACAAGGAAAGAACGGACGGTATCGTTCACTTATAACCAATACGCTCAGATGATGAACATCAGTAGGGAAAATGCTTATGGTGTATTAGCTAAAGCAACCAGAGAGCTGATGACGAGAACTGTGGAAATCAGGAATCCTTTGGTTAAAGGCTTTGAGATTTTCCAGTGGACAAACTATGCCAAGTTCTCAAGCGAAAAATTAGAATTAGTTTTTAGTGAAGAGATATTGCCTTATCTTTTCCAGTTAAAAAAATTCATAAAATATAATCTGGAACATGTTAAGTCTTTTGAAAACAAATACTCTATGAGGATTTATGAGTGGTTATTAAAAGAACTAACACAAAAGAAAACTCACAAGGCAAATATAGAGATTAGCCTTGATGAATTTAAGTTCATGTTAATGCTTGAAAATAACTACCATGAGTTTAAAAGGCTTAACCAATGGGTTTTGAAACCAATAAGTAAAGATTTAAACACTTACAGCAATATGAAATTGGTGGTTGATAAGCGAGGCCGCCCGACTGATACGTTGATTTTCCAAGTTGAACTAGATAGACAAATGGATCTCGTAACCGAACTTGAGAACAACCAGATAAAAATGAATGG TGACAAAATACCAACAACCATTACATCAGATTCCTACCTACGTAACGGACTAAGAAAAACACTACACGATGCTTTAACTGCAAAAATTCAGCTCACCAGTTTTGAGGCAAAATTTTTGAGTGACATGCAAAGTAAGCATGATCTCAATGGTTCGTTCTCATGGCTCACGCAAAAACAACGAACCACACTAGAGAACATACTGGCTAAATACGGAAGGATCTGAGGTTCTTATGGCTCTTGTATCTATCAGTGAAGCATCAAGACTAACAAACAAAAGTAGAACAACTGTTCACCGTTAGATATCAAAGGGAAAACTGTCCATATGCACAGATGAAAACGGTGTAAAAAAGATAGATACATCAGAGCTTTTACGAGTTTTTGGTGCATTTAAAGCTGTTCACCATGAACAGATCGACAATGTAACTACCTCCTTCGTTGAGAACTCACAATTATATTACCAATGCTTAATCAGTGAGGCACCTATCTCAGCGATCTGTCTATTTCGTTCATCCATAGTTGCCTGACTCCCCGTCGTGTAGATAACTACGATACGGGAGGGCTTACCATCTGGCCCCAGTGCTGCAATGATACCGCGAGACCCACGCTCACCGGCTCCAGATTTATCAGCAATAAACCAGCCAGCCGGAAGGGCCGAGCGCAGAAGTGGTCCTGCAACTTTATCCGCCTCCATCCAGTCTATTAATTGTTGCCGGGAAGCTAGAGTAAGTAGTTCGCCAGTTAATAGTTTGCGCAACGTTGTTGCCATTGCTACAGGCATCGTGGTGTCACGCTCGTCGTTTGGTATGGCTTCATTCAGCTCCGGTTCCCAACGATCAAGGCGAGTTACATGATCCCCCATGTTGTGCAAAAAAGCGGTTAGCTCCTTCGGTCCTCCGATCGTTGTCAGAAGTAAGTTGGCCGCAGTGTTATCACTCATGGTTATGGCAGCACTGCATAATTCTCTTACTGTCATGCCATCCGTAAGATGCTTTTCTGTGACTGGTGAGTACTCAACCAAGTCATTCTGAGAATAGTGTATGCGGCGACCGAGTTGCTCTTGCCCGGCGTCAATACGGGATAATACCGCGCCACATAGCAGAACTTTAAAAGTGCTCATCATTGGAAAACGTTCTTCGGGGCGAAAACTCTCAAGGATCTTACCGCTGTTGAGATCCAGTTCGATGTAACCCACTCGTGCACCCAACTGATCTTCAGCATCTTTTACTTTCACCAGCGTTTCTGGGTGAGCAAAAACAGGAAGGCAAAATGCCGCAAAAAAGGGAATA AGGGCGACACGGAAATGTTGAATACTCATACTCTTCCTTTTTCAATATTATTGAAGCATTTATCAGGGTTATTGTCTCATGAGCGGATACATATTTGAATGTATTTAGAAAAATAAACAAATAGGGGTTCCGCGCACATTTCCCCGAAAAGTGCCACCTGACGTCTAAGAAACCATTATTATCATGACATTAACCTATAAAAATAGGCGTATCACGAGGCCCTTTCGTCTCGCGCGTTTCGGTGATGACGGTGAAAACCTCTGACACATGCAGCTCCCGGAGACGGTCACAGCTTGTCAACAAGACCTGACCTAACGGTAAGAttattaagcaccggtggagtgacgaccttcagcacgttcgtactgttcaacgatggtgtagtcttcgttgtgggaggtgatgtccagtttgatgtcggttttgtaagcacccggcagctgaaccggttttttagccatgtaggtggttttaacttcagcgtcgtagtgaccaccgtctttcagtttcagacgcattttgatttcacctttcagagcaccgtcttccgggtacatacgttcggtggaagcttcccaacccatggtttttttctgcataaccggaccgtcggacgggaagttggtaccacgcagtttaactttgtagatgaactcaccgtcttgcagggaggagtcctgggtaacggtaacaacaccaccgtcttcgaagttcataacacgttcccatttgaaaccttccgggaaggacagtttcaggtagtccgggatgtcagccgggtgtttaacgtaagctttggaaccgtactggaactgcggggacaggatgtcccaagcgaacggcagcggaccacctttggtaactttcagtttagcggtctgggtaccttcgtacggacgaccttcaccttcaccttcgatttcgaactcgtgaccgttaacggaaccttccatacgaactttgaaacgcatgaactctttgataacgtcttcggaggaagccatctagtactttcctgtgtgactctagtagctagcactgtacctaggactgagctagccgtcaaACGATTGGTAAACCCGGTGaacgcatgagAAAGCCCCCGGAAGATCACCTTCCGGGGGCTTTtttattgcgcGGACCAAAACGAAAAAAGACGCTCGAAAGCGTCTCTTTTCTGGAATTTGGTACCGAGCATTCGATCAGCCAAACGTCTCTTCAGGCCACTGACTAGCGATAACTTTCCCCACAACGGAACAACTCTCATTGCATGGGATCATTGGGTACTGTGGGTTTAGTGGTTGTAAAAACACCTGACCGCTATCCCTGATCAGTTTCTTGAAGGTAAACTCATCACCCCCAAGTCTGGCTATGCAGAAATCACCTGGCTCAACAGCCTGCTCAGGGTCAACGAGAATTAACATTCCGTCAGGAAAGCTTGGCTTGGAGCCTGTTGGTGCGGTCATGGAATTACCTTCAACCTCAAGCCAGAATGCAGAATCACTGGCTTTTTTGGTTGTGCTTACCCATCTCTCCGCATCACCTTTGGTAAAGGTTCTAAGCTTAGGTGAGAACATCCCTGCCTGAACAT GAGAAAAAACAGGGTACTCATACTCACTTCTAAGTGACGGCTGCATACTAACCGCTTCATACATCTCGTAGATTTCTCTGGCGATTGAAGGGCTAAATTCTTCAACGCTAACTTTGAGAATTTTTGTAAGCAATGCGGCGTTATAAGCATTTAATGCATTGATGCCATTAAATAAAGCACCAACGCCTGACTGCCCCATCCCCATCTTGTCTGCGACAGATTCCTGGGATAAGCCAAGTTCATTTTTCTTTTTTTCATAAATTGCTTTAAGGCGACGTGCGTCCTCAAGCTGCTCTTGTGTTAATGGTTTCTTTTTTGTGCTCATACGTTAAATCTATCACCGCAAGGGATAAATATCTAACACCGTGCGTGTTGACTATTTTACCTCTGGCGGTGATAATGGTTGCATagctgtcaccggatgtgctttccggtctgatgagtccgtgaggacgaaacagcctctacaaataattttgtttaaAACAACACCCACTAAGATAAGGTAGAAACATGAGCAAAGGAGAAGAACTTTTCACTGGAGTTGTCCCAATTCTTGTTGAATTAGATGGTGATGTTAATGGGCACAAATTTTCTGTCCGTGGAGAGGGTGAAGGTGATGCTACAAACGGAAAACTCACCCTTAAATTTATTTGCACTACTGGAAAACTACCTGTTCCGTGGCCAACACTTGTCACTACTCTGACCTATGGTGTTCAATGCTTTTCCCGTTATCCGGATCACATGAAACGGCATGACTTTTTCAAGAGTGCCATGCCCGAAGGTTATGTACAGGAACGCACTATATCTTTCAAAGATGACGGGACCTACAAGACGCGTGCTGAAGTCAAGTTTGAAGGTGATACCCTTGTTAATCGTATCGAGTTAAAGGGTATTGATTTTAAAGAAGATGGAAACATTCTTGGACACAAACTTGAGTACAACTTTAACTCACACAATGTATACATCACGGCAGACAAACAAAAGAATGGAATCAAAGCTAACTTCAAAATTCGCCACAACGTTGAAGATGGTTCCGTTCAACTAGCAGACCATTATCAACAAAATACTCCAATTGGCGATGGCCCTGTCCTTTTACCAGACAACCATTACCTGTCGACACAATCTGTCCTTTCGAAAGATCCCAACGAAAAGCGTGACCACATGGTCCTTCTTGAGTTTGTAACTGCTGCTGGGATTACACATGGCATGGATGAGCTCTACAAATGAgttggtctggtgtcaaaaataaCTCaatacgGTCGACagttaaccaaAAAGGGGGGATTTTATCTCCCCTTTaatttttcctGTCTCCCcctttggtcgAAAAAAAAAGCCCGC ACTGTCAGGTGCGGGCTTTTTTctgtgtttccCTTGAAGTAATGTATACGACAGAGTCCGTGCACCTACCAA.

### Bacterial genome sequencing

Bacterial genomic DNA from each strain was purified individually using the QIAamp Fast DNA Stool Mini Kit (Qiagen, #51604). DNA was quantified by Nanodrop. 250 ng of input DNA was prepared for sequencing using the NEBNext Ultra II DNA Library Prep Kit (New England Biolabs, #E7645S) and sequencing adaptors were ligated using NEBNext Multiplex Oligos for Illumina (#E7500S, #E7335S) as described. Bacterial DNA was amplified by using 4 cycles according to the NEBNext protocol. Libraries were quantified using a Library Quantification Kit (Kapa Biosystems, #KK4824) and sequenced on a NextSeq550 High Output. Deep sequencing reads were then aligned to the CP022686.1 reference genome for E. Coli Nissle 1917.

### D-LA production kinetics

To validate D-LA-producing bacteria *in vivo*, control bacteria and EcN^Lac^ were administrated by oral gavage for a week. 1, 4 and 24 hours after the last gavage, plasma was collected in heparinized tubes and fecal content from small intestine and colon was collected in BeadBug prefilled tubes (Millipore Sigma, #Z763829), weighted and diluted in 0.5 mL of cold and sterile 1X PBS. Next, samples were homogenized in PowerLyzer 24 Homogenizer (Qiagen, #13155) two times at 4m/s for 10 seconds, and then, at maximal speed for 20 min at 4°C. Supernatants were used to quantify Colony-forming Units (CFU), D-lactate and L-lactate levels. In other set of experiments, small intestine and colon were isolated 1, 4 and 24 h after D-LA bacteria gavage and digested for flow cytometry analysis. To evaluate EcN^Lac^ protective role, control or EcN^Lac^ bacteria were gavaged every day starting 3 days before EAE induction or weekly starting the day of EAE induction.

### D-lactate and L-lactate quantification

For D-lactate, feces and plasma were filtered on 10kD Spin Column (Abcam, #ab93349) to concentrate D-lactate. For L-lactate, samples were deproteinized by PEG precipitation. D-and L-lactate concentrations were quantified using D-Lactate Assay Kit (Abcam, #ab174096) and L-lactate Assay Kit (Biomedical Research, Service and Clinical Application #A-108) as indicated in the manufacturer’s instructions.

### CFUs

Feces supernatants were serial diluted and seeded in LB Agar media with 300 µg/mL of streptomycin for control bacteria (Syn094) or 100 µg/mL of carbenicillin/ampicillin for D-LA-producing bacteria (Syn028). Next day, formed colonies were counted the dilution factor was corrected.

### Inducible activation analysis in vitro

D-LA-producing bacteria or GFP^+^ bacteria (SYNB2189) were diluted in 2x YT medium (Sigma Aldrich, #Y1003-500ML) with 100 µg/mL of carbenicillin/ampicillin or 100 µg/mL of carbenicillin, respectively and shacked at 250 rpm at 37C. At indicated time points, D-LA was quantified as previously described and GFP expression was measured by FACs on a Symphony A5 (BD Biosciences).

### Mouse DC sorting

cDCs from CNS and spleen were sorted as B220/Ly6G/Ly6C/TER119/O4^neg^;CD45^high^CD11c^high^MHCII^high^. Compensation was performed on single-stained beads. Cells were sorted on a FACS Aria Ilu (BD Biosciences).

### Bulk RNA-sequencing

Sorted cells were lysed on extraction buffer and RNA was isolated using the PicoPure RNA Isolation Kit (Thermo Fisher Scientific, #KIT0204). RNA was suspended in 10 µL of nuclease free water at a concentration of 0.5-1 ng/µL and sequenced using or SMARTSeq2^76^ at the Broad Institute. All bulk RNA-sequencing results were aligned to EMSEMBL GRCm38.p6 reference genome and quantified using Kallisto (v0.46.1)^77^ and Salmon (v 1.1.0)^78^. DESeq2 software was used to perform differential expression analysis and the apeGLM algorithm^79^ was used to remove noise in log2 fold change analyses following differential expression analysis.

### Pathway analysis

GSEA or GSEAPreranked analyses were used to generate enrichment plots for bulk RNA-seq or scRNA-seq data^80, 81^ using MSigDB molecular signatures for canonical pathways: KEGG/Reactome/Biocarta (c2.cp.all), Motif (c3.all), Gene ontology (c5.cp.all), and Hallmark (h.all). To determine regulators of gene expression networks, Ingenuity Pathway Analysis software (Qiagen) was used by inputting gene expression datasets with corresponding log (FoldChange) expression levels compared to other groups. “Canonical pathways” and “upstream analysis” metrics were considered significant at p<0.05.

### Tissue Immunofluorescence

For immunofluorescence, distal ileum and distal colon were dissected after one week of EcN or EcN^Lac^ treatment, then fixed in 4% paraformaldehyde for 4hrs at 4°C. Tissue was subsequently dehydrated by incubating in 10% sucrose in PBS for 1hr at 4°C, then 20% sucrose in PBS for 1hr at 4°C, then 30% sucrose overnight at 4°C. Tissues were frozen in OCT (Sakura, #4583) and 8 µm sections were prepared by cryostat on SuperFrost Plus Gold slides (Fisher Scientific, #15-188-48). Sections were permeabilized with 1X permeabilization buffer (BD Biosciences, #554723) for 10 minutes at RT, then blocked using serum-free protein block (Agilent, #X0909) for 10 minutes at RT. Sections were then incubated with primary antibodies diluted in 1X permeabilization buffer overnight at 4°C. Following primary antibody incubation, sections were washed 3X with 1X permeabilization buffer and incubated with secondary or conjugated antibodies diluted in 1X permeabilization buffer for 1 hour at RT. After secondary and conjugated antibody incubation, sections were washed 3X with 1X permeabilization buffer and mounted with ProLong Gold Antifade Mountant (Fisher Scientific, #P36930). Primary and conjugated antibodies used in this study were: Armenian hamster anti-CD11c pacific blue (Biolegend, 1:100, #117321), goat anti-HIF-1α (R&D Systems, 1:100, AF1935-SP), rabbit anti-NDUFA4L2 (Proteintech, 1:50, 16480-1-AP) and mouse anti-sXBP1 (Biolegend, 1:50, 647502). Secondary antibodies used in this study were: donkey anti-goat IgG Alexa Fluor 488 (Jackson Immunoresearch, #705-545-003), donkey anti-rabbit IgG (H+L) Highly Cross-Adsorbed, Alexa Fluor 568 (Life Technologies, #A10042), and donkey anti-mouse IgG H&L, Alexa Fluor 647 (Abcam, ab150107), all at 1:1000 working dilution. Sections were imaged and tile scan images were stitched on an LSM-880-AiryScan confocal microscope using the ZEN Black software (Zeiss), and image analysis was performed with FIJI version of ImageJ (NIH). DCs expression of Hif1⍺, NDUFA4L2, and sXBP1 in colon and small intestines sections was quantified as previously described (PMID: 34294479, 33712740). Briefly, CD11c-positive cells were detected by implementing the bwlabel algorithm for labeling connected components in 2-D images (Book chapter reference: Haralick, Robert M., and Linda G. Shapiro, Computer and Robot Vision, Volume I, Addison-Wesley, 1992, pp. 28-48.) using MATLAB (MathWorks). Small puncta sized < 50 μm2 were regarded as artifacts and were filtered out. Blood vessel related fluorescence artifacts were manually removed. Next, the averaged Hif1⍺, NDUFA4L2, and sXBP1 fluorescence intensities were calculated per individual CD11c-positive cells and outliers were removed using the Robust Regression and Outlier Removal (ROUT) method with coefficient Q = 1% (PMID: 16526949).

### Data deposition

All raw and processed deep sequencing data have been deposited into GEO.

---

**Extended Data Figure 1.**
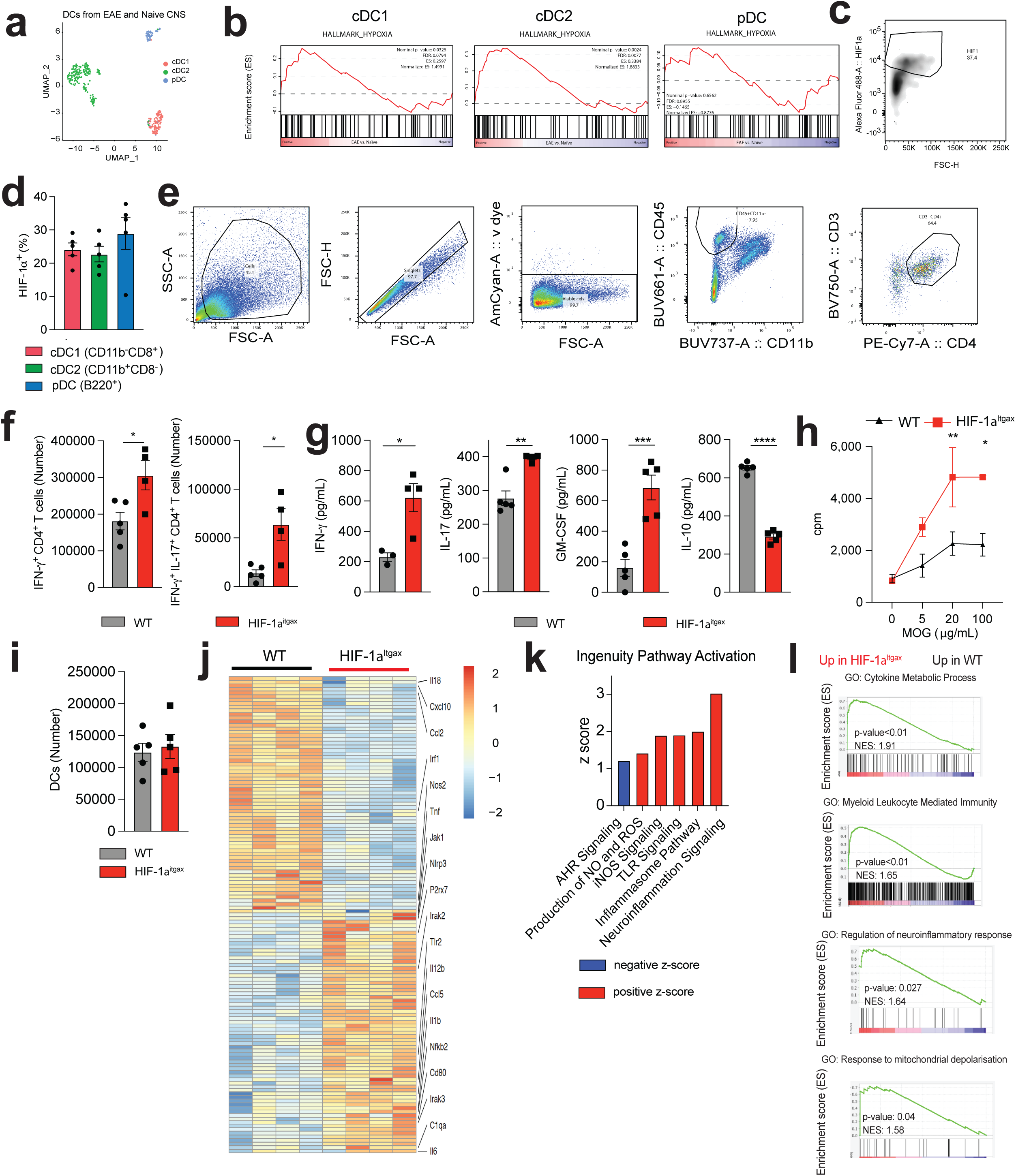
Analysis of HIF-1^Itgax^ mice during EAE. **(a)** Uniform Manifold Approximation and Projection (UMAP) displaying CNS DCs analyzed by scRNAseq during EAE. **(b)** GSEA of hypoxia activation in cDC1, cDC2 and pDC from scRNAseq dataset^71^. **(c,d)** Representative dot plot (c) and flow cytometry analysis (d) of HIF-1α expression in splenic cDC1s (CD8^+^CD11b^-^), cDC2s (CD8^-^CD11b^-^) and pDCs (B220^+^) 25 days after EAE induction. **(e)** Gating strategy used to analyze CD4^+^ T cells in the CNS. **(f)** IFN-γ^+^, IFN-γ^+^IL-17^+^ CD4^+^ T cells in spleen of WT and HIF-1α^Itgax^ mice. **(g,h)** Cytokine (g) and proliferative (g) splenocyte recall response to 20 μg/mL of MOG35-55 . **(i)** DCs in CNS from WT and HIF-1α^Itgax^ mice. **(j-l)** Heatmap (j), IPA (k) and GSEA (l) from RNAseq of CNS DCs in WT or HIF-1α^Itgax^. Data shown as mean±SEM. ****p<0.0001, *p<0.05, ns: p>0.05.

**Extended Data Figure 2.**
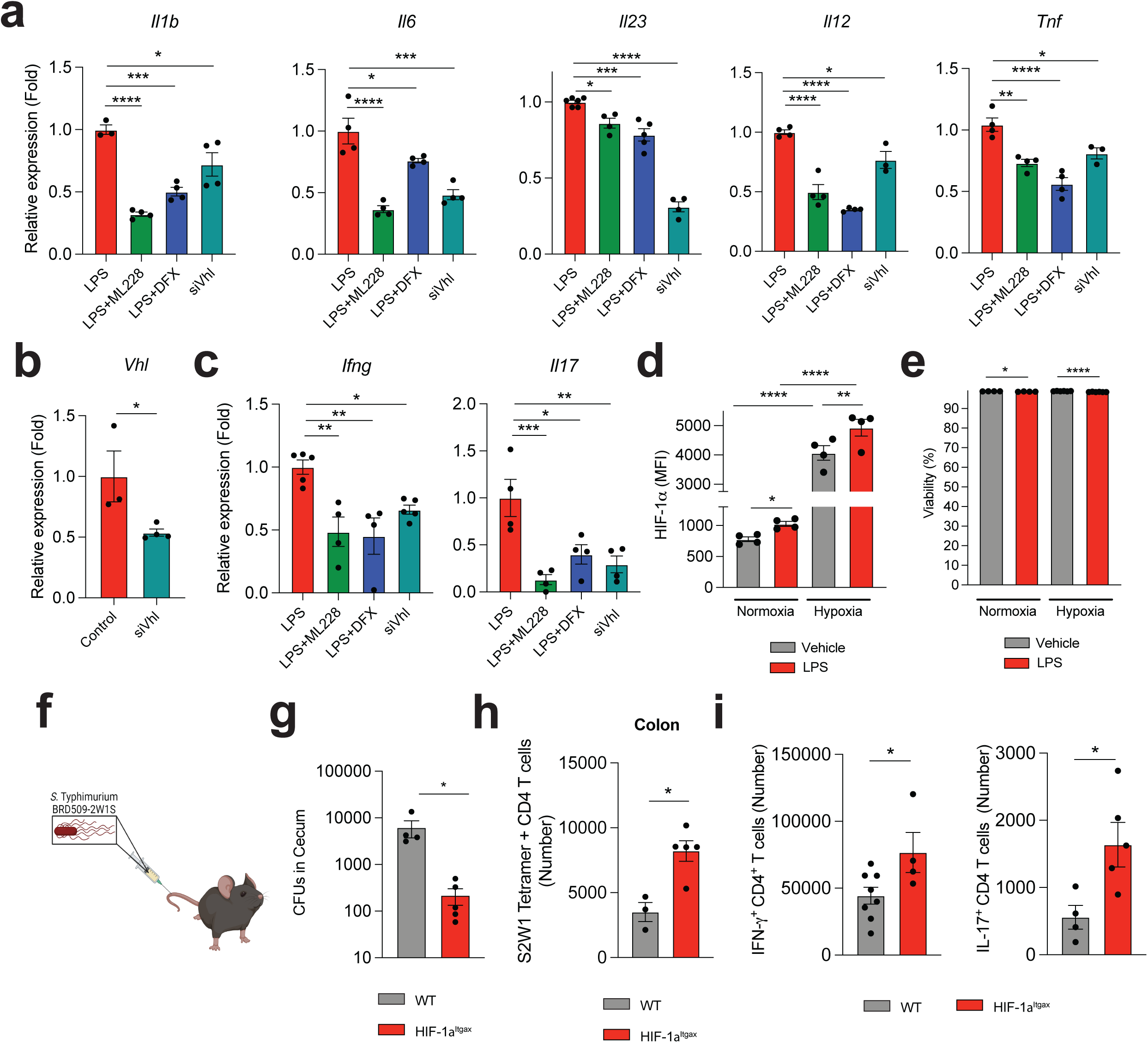
Effects of HIF-1α in DCs. **(a)** mRNA expression in DCs treated with LPS, ML228 or DFX, or after *Vhl* knockdown (si*Vhl*). **(b)** *Vhl* expression in si*Vhl-* treated BMDCs. **(c)** *Ifng* and *Il17* expression in 2D2^+^ CD4^+^ T cells co-cultured with WT BMDCs pre-stimulated with ML228, DFX or si*Vhl* and LPS. **(d,e)** HIF-1α MFI (a) and frequency of viable cells (b) following LPS stimulation under hypoxia and normoxia. **(f,g)** Experimental design (f) and *S. thyphimurium* CFU quantification (g) in cecum from HIF-1α^Itgax^ and WT mice 14 days after infection. **(h,i)** S2W1 Tetramer-specific (h) and IFN-γ^+^ and IL-17^+^ (i) CD4^+^ T cells in colon from HIF-1α^Itgax^ and WT mice 14 days after S. thyphimurium infection. Data shown as mean±SEM. ****p<0.0001, **p<0.01, *p<0.05, ns: p>0.05.

**Extended Data Figure 3.**
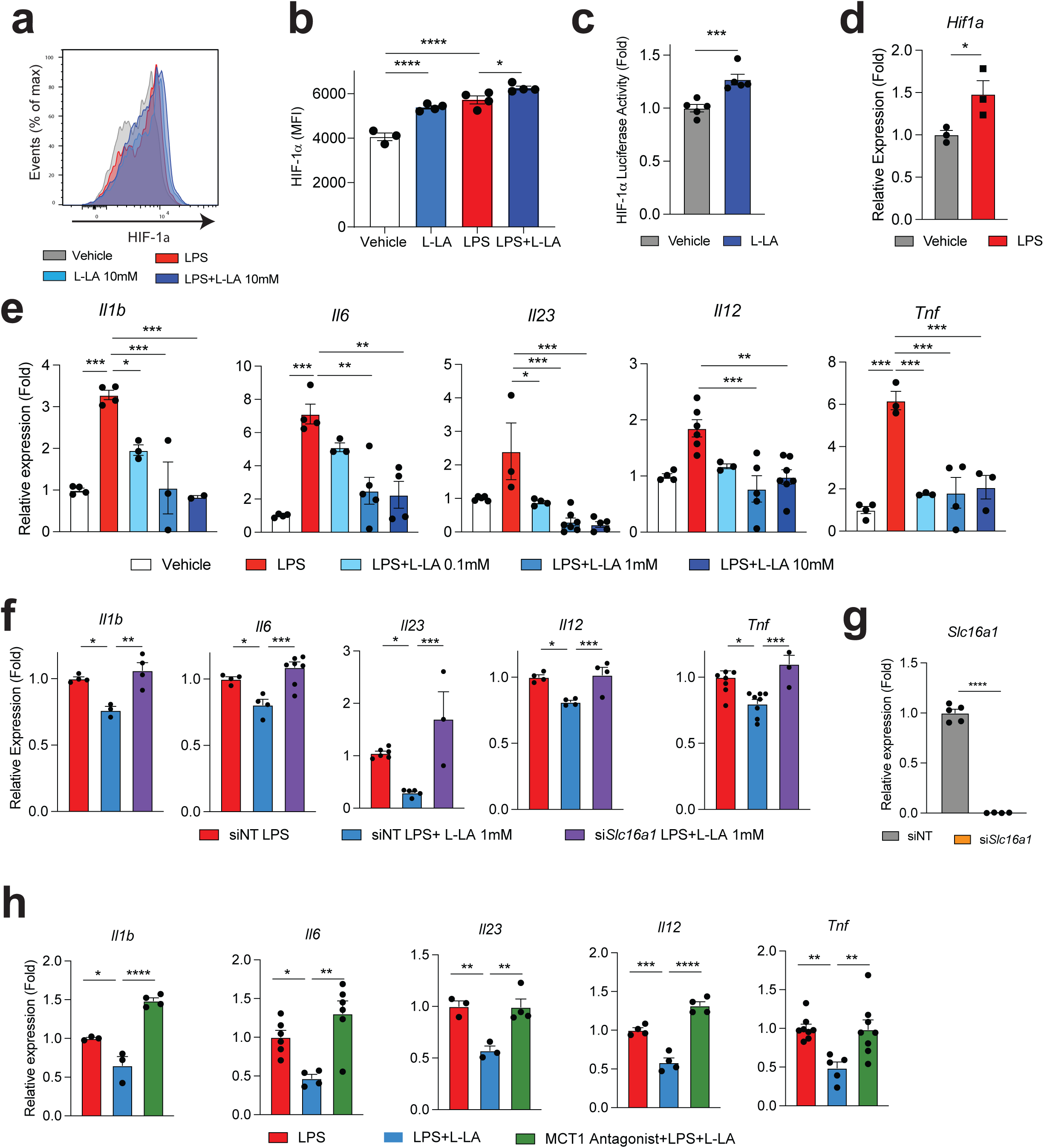
Effects HIF-1α activation by L-LA on DCs. **(a,b)** Representative histogram (a) and MFI (b) of HIF-1α expression in mouse WT BMDCs treated with LPS and L-LA. **(c)** HIF-1α luciferase activity in FVB.129S6-Gt(ROSA)26Sor^tm2(HIF1A/luc)Kael/J^ BMDCs treated with LA. **(d)** *Hif1a* expression in LPS treated BMDCs. **(e)** mRNA expression in WT BMDCs treated with L-LA. **(f-h)** mRNA expression in WT BMDCs after *Slc16a1* knockdown (f,g) or treatment with the MCT1-antagonist AZD3965 (h). Data shown as mean±SEM. ****p<0.0001, **p<0.01, *p<0.05, ns: p>0.05.

**Extended Data Figure 4.**
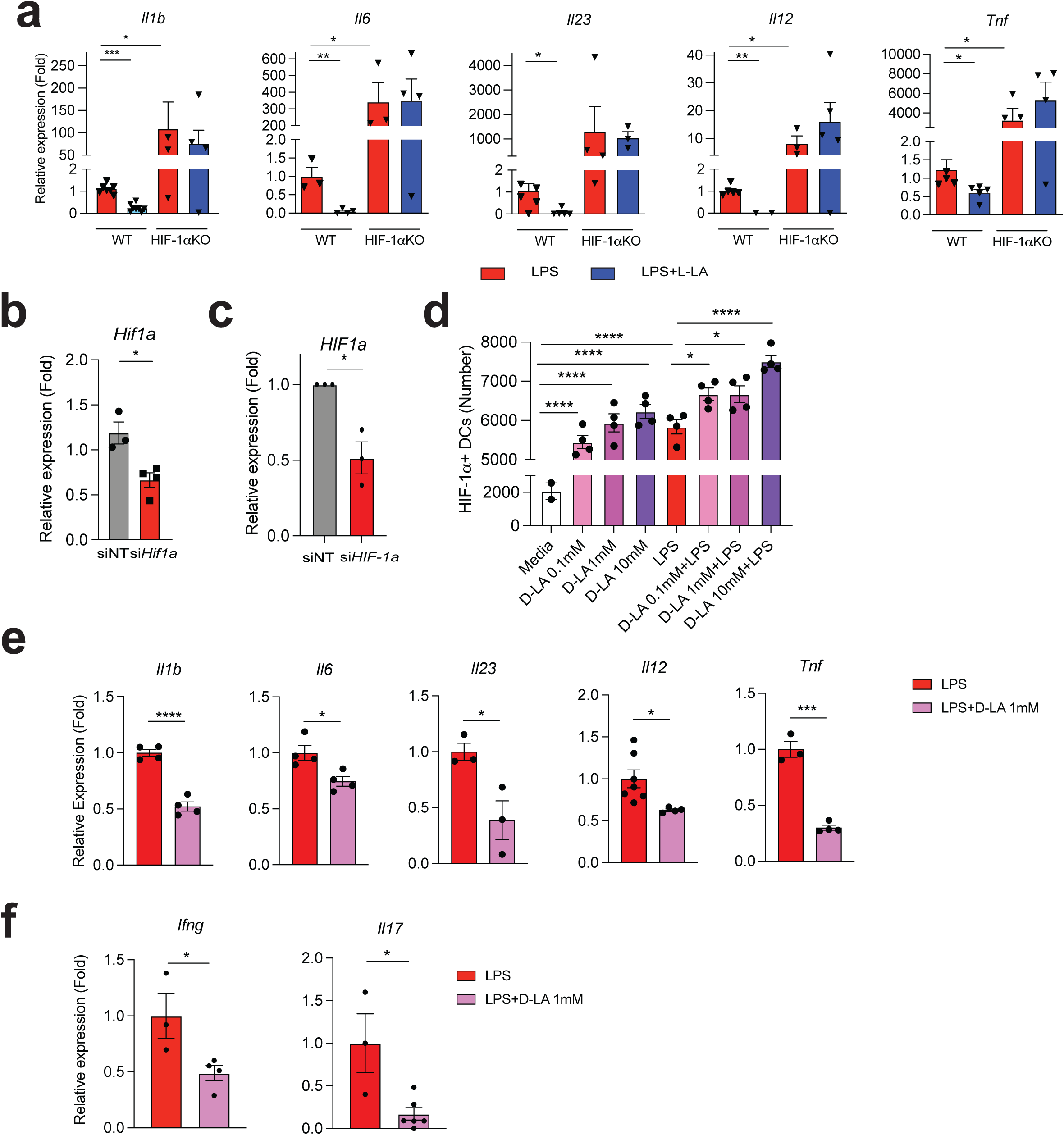
Effect of HIF-1α activation by D-LA on DCs. **(a)** mRNA expression in HIF-1α^Itgax^ or WT splenic DCs stimulated with LPS or L-LA. **(b,c)** Knockdown of *Hif1a* (b) and *HIF1a* (c) in mouse BMDCs (b) and human DCs (c). **(d)** HIF-1α^+^ DCs following treatment with D-LA and LPS. **(e)** mRNA expression in WT splenic DCs after LPS or D-LA (1mM) treatment for 6h. **(f)** *Ifng* and *Il17* expression in 2D2 CD4^+^ T cells co-cultured with LPS or D-LA (1mM) pre-treated DCs. Data shown as mean±SEM. ****p<0.0001, **p<0.01, *p<0.05, ns: p>0.05.

**Extended Data Figure 5.**
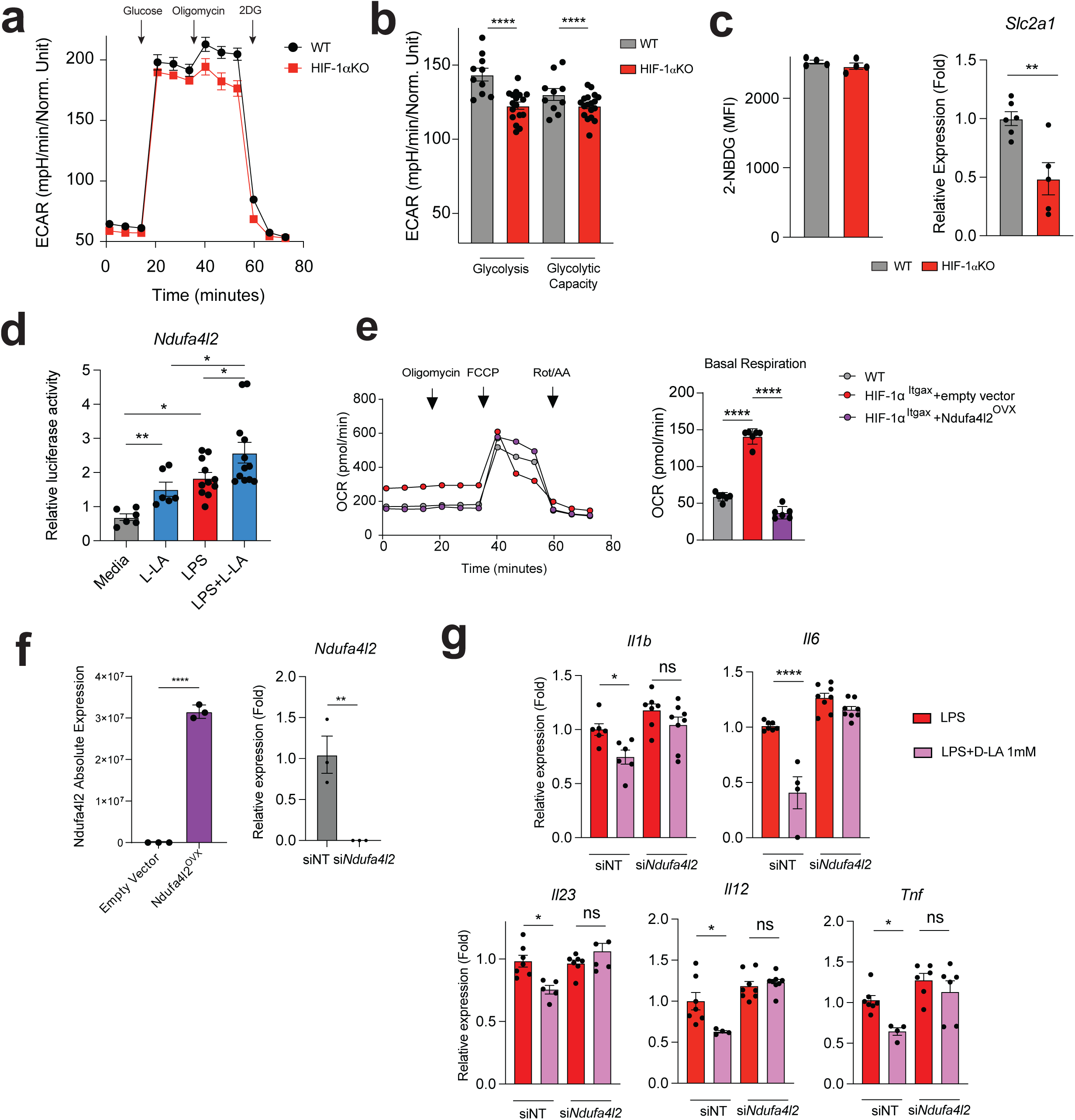
Effect of HIF-1α on DC metabolism. **(a)** ECAR in WT and HIF-1α^Itgax^ BMDCs after glucose, oligomycin and 2-DG treatment. **(b)** Glycolysis and glycolytic capacity in WT and HIF-1α^Itgax^ BMDCs. **(c)** 2-NBDG uptake and *Slc2a1* expression in WT and HIF-1α^Itgax^ BMDCs after LPS stimulation. **(d)** Transactivation of *Ndufa4l2* promoter in Ndufa4l2-luciferase transfected DC2.4 cells treated with L-LA or LPS for 24h. **(e)** OCR in WT or HIF-1α^Itgax^ BMDCs overexpressing *Ndufa4l2*. **(f)** *Ndufa4l2* expression after transfection with Ndufa4l2-oxerexpression plasmid or silencing with siRNA. **(g)** mRNA expression in Ndufa4l2-silenced DCs treated with LPS or D-LA for 6h. Data shown as mean±SEM. ****p<0.0001, **p<0.01, *p<0.05, ns: p>0.05.

**Extended Data Figure 6.**
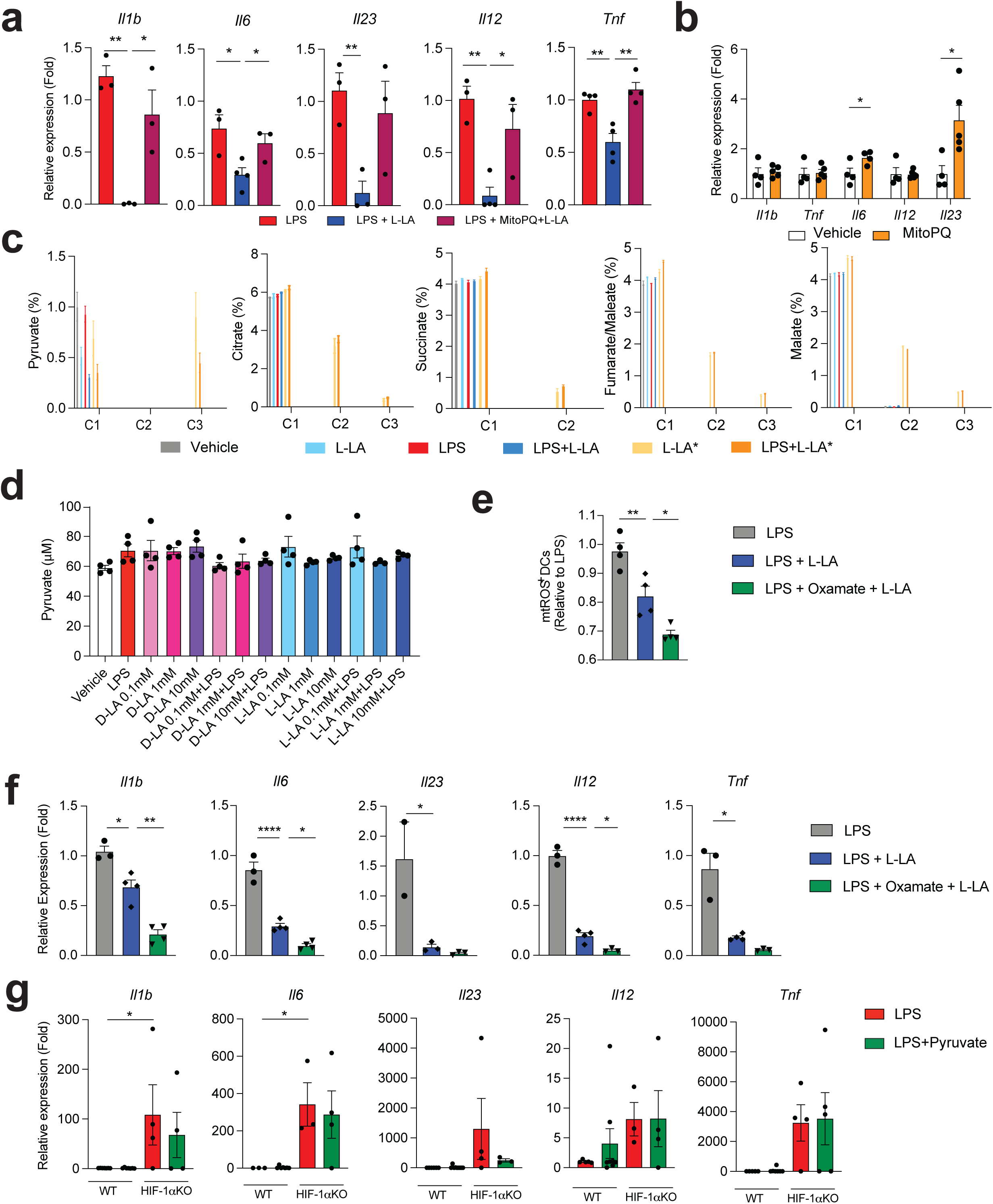
L-LA incorporation to TCA intermediates. **(a)** mRNA expression in BMDCs stimulated with LPS, MitoPQ or L-LA for 6h. **(b)** mRNA expression in DCs treated with mitoPQ for 6h. **(c)** 13C incorporation into TCA intermediates in BMDCs treated with uniformly labeled 13C lactate (L-LA*), L-LA or LPS for 1h. **(d)** Pyruvate intracellular levels in DCs after treatment with L-LA or LPS for 1h. **(e,f)** mtROS production (e) and mRNA expression (f) in DCs pre-treated with LDH-inhibitor oxamate, L-LA or LPS. **(g)** mRNA expression in splenic WT and HIF-1α^Itgax^ DCs treated with LPS or pyruvate for 6h. Data shown as mean±SEM. ****p<0.0001, **p<0.01, *p<0.05, ns: p>0.05.

**Extended Data Figure 7.**
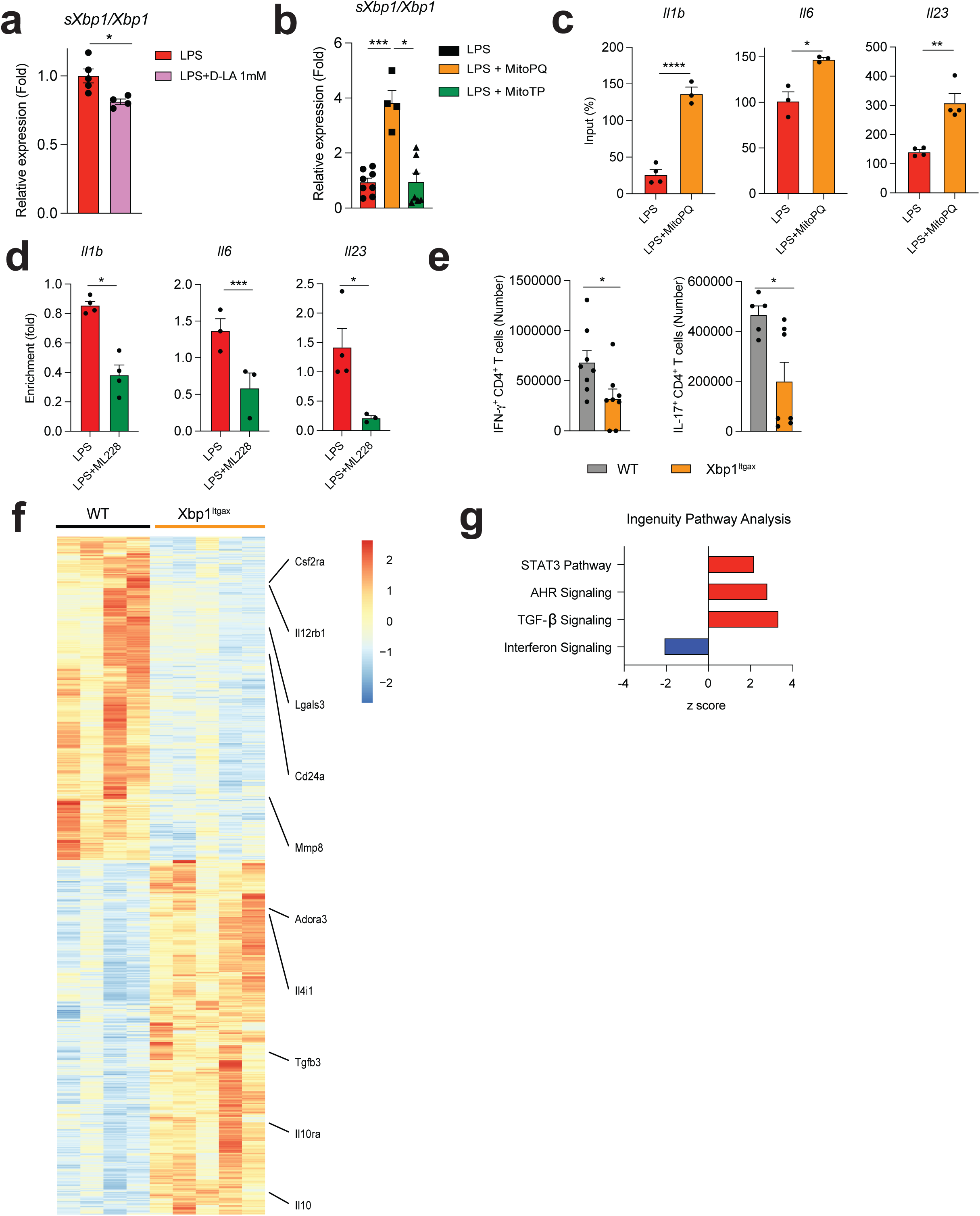
Control of DCs by XBP1. **(a,b)** *sXbp1/Xbp1* in BMDCs treated with LPS or LPS+D-LA (a) and LPS, MitoPQ or MitoTempo (MitoTP) (b) for 6h. **(c,d)** XBP1 recruitment to *Il1b*, *Il6* and *Il23* promoters in BMDCs treated with LPS and mitoPQ (c) or LPS and ML228 (d) for 6h. **(e)** IFNγ^+^ and IL-17^+^ CD4 splenic T cells in WT and Xbp1^Itgax^ mice 30 days after EAE induction. **(f,g)** Heatmap (f) and IPA (g) in CNS DCs from WT and Xbp1^Itgax^ mice. Data shown as mean±SEM. ***p<0.001, **p<0.01, *p<0.05, ns: p>0.05.

**Extended Data Figure 8.**
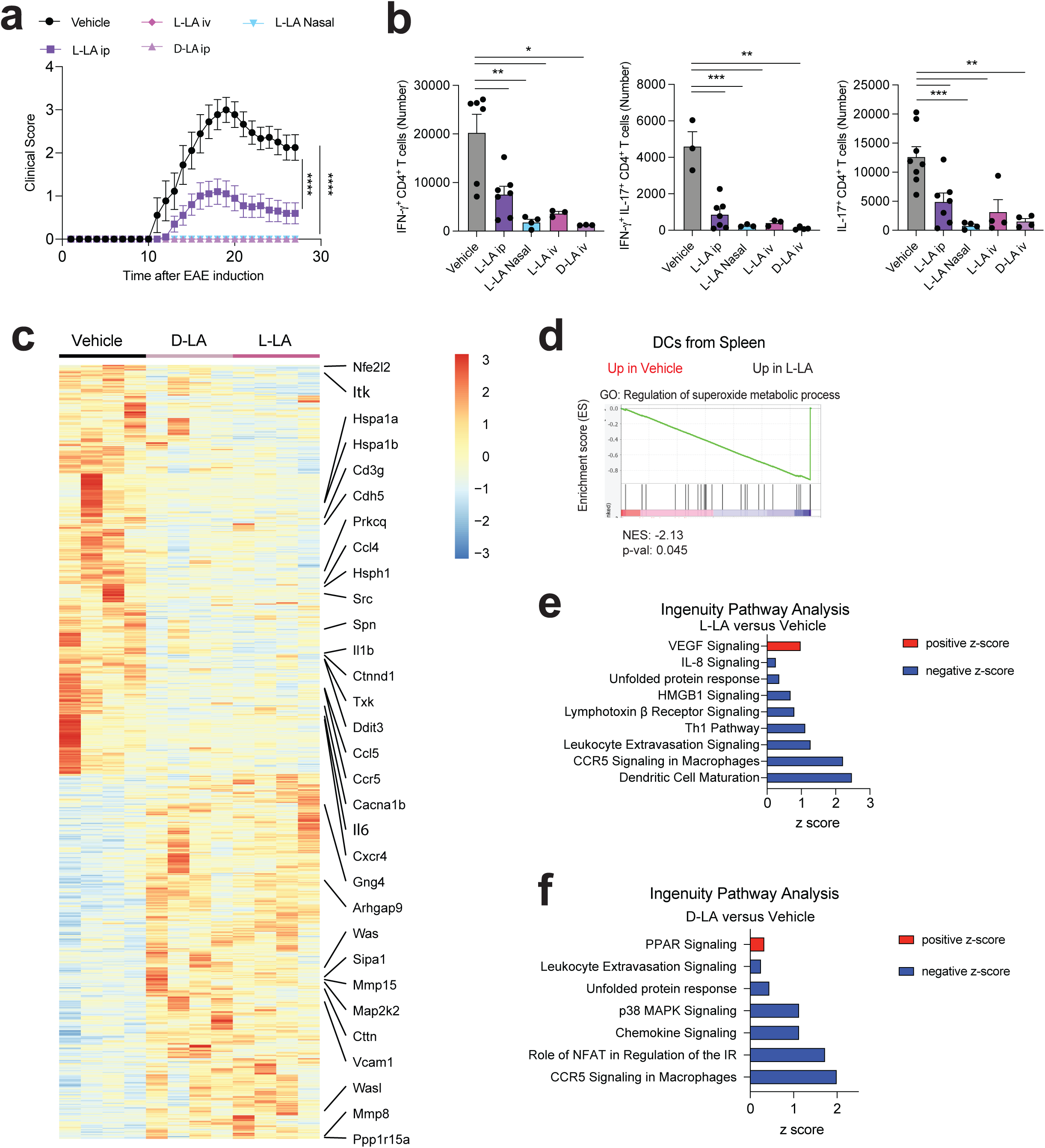
Effect of LA on EAE. **(a,b)** EAE development (a) and IFN-γ^+^, IFN-γ^+^ IL-17^+^ and IL-17^+^ CD4^+^ T cells in CNS (b) after intraperitoneal (ip), nasal or intravenous (iv) L-LA or D-LA administration. **(c-f)** Heatmap (c), GSEA (d) and IPA in splenic DCs from mice treated with L-LA(e) and D-LA (f) analyzed 28 days after EAE induction. Data shown as mean±SEM. ***p<0.001, **p<0.01, *p<0.05, ns: p>0.05.

**Extended Data Figure 9.**
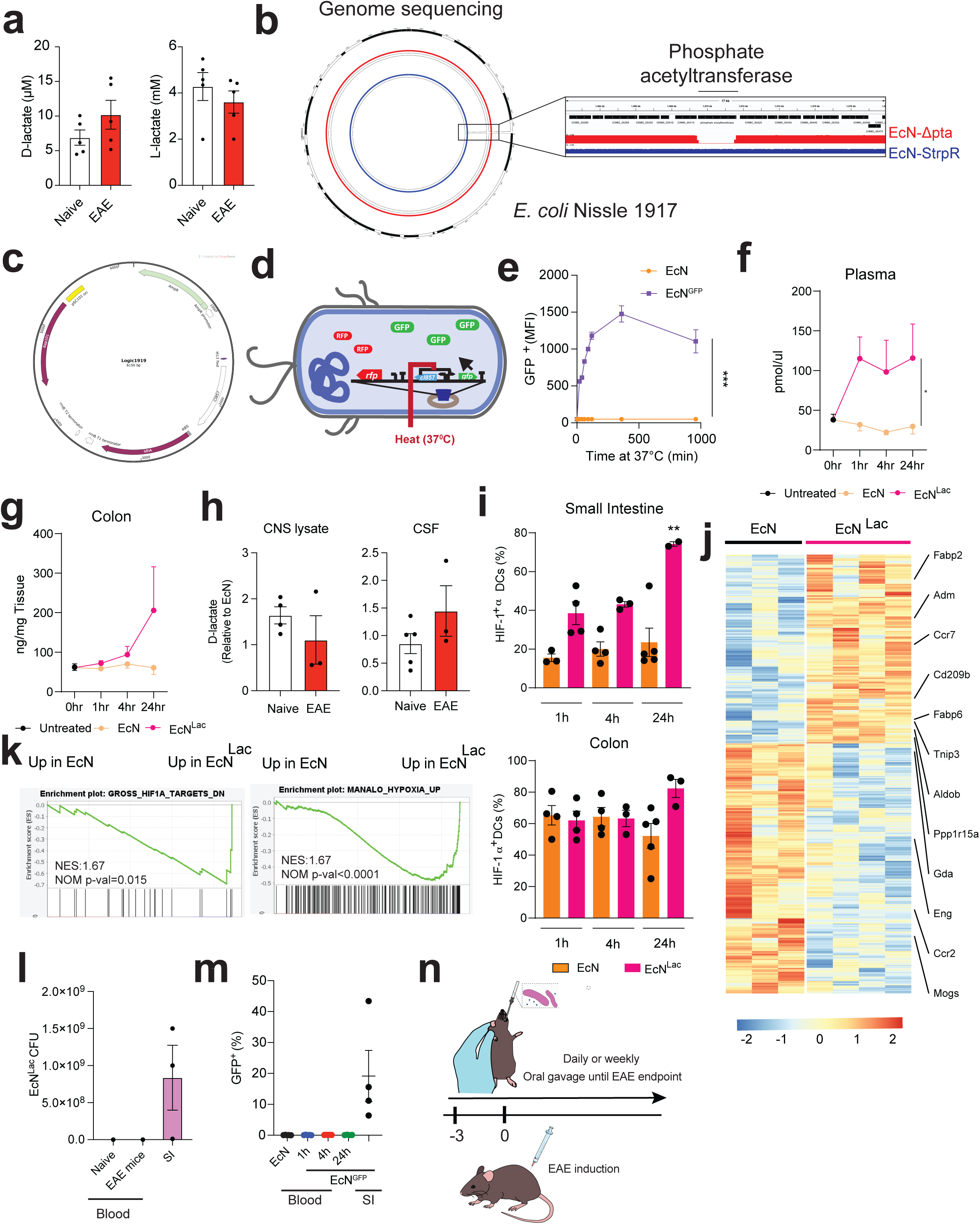
EcN^Lac^ characterization. **(a)** L-LA and D-LA in plasma from naïve and peak EAE mice. **(b)** Genome sequencing of EcN^Lac^ and parental EcN strains. **(c)** Plasmid used for *ldhA* expression. **(d)** EcN^GFP^ reporter strain. **(e)** GFP expression in EcN^GFP^ after activation at 37C. **(f,g)** D-LA in plasma (e) and colon tissue (f) after EcN^Lac^ or EcN administration. **(h)** D-LA in CSF and CNS lysate from naïve and EAE mice after EcN^Lac^ administration, shown relative to D-LA in EcN-treated mice. **(i)** Small intestine and colon HIF-1α^+^ DCs in EcN-or EcN^Lac^-treated mice. **(j,k)** heatmap (j) and GSEA (k) of small intestine DCs from EcN-or EcN^Lac^-treated mice analyzed by RNA-seq. **(l, m)** EcN^Lac^ in blood from naïve and EAE mice treated with EcN^Lac^ for a week (l) and EcN^GFP^ in blood 1, 4 and 24h after oral gavage (m). Small intestine content is shown as a positive control. **(n)** Experimental design. Data shown as mean±SEM. **p<0.01, *p<0.05, ns: p>0.05.

**Extended Data Figure 10.**
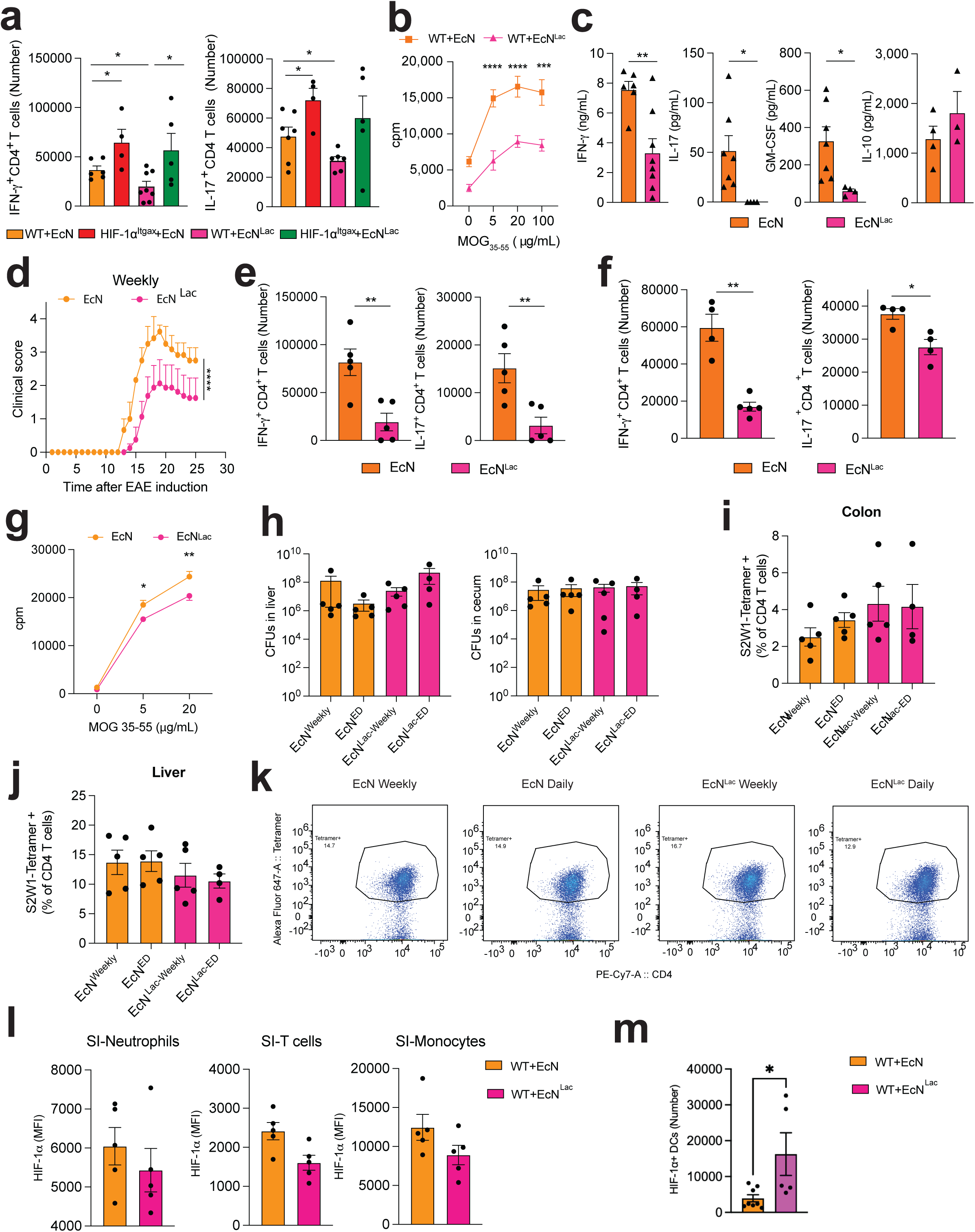
Effects of EcN^Lac^ on EAE and *S. thyphimurium* infection. **(a-c)** Splenic IFN-γ^+^ and IL-17^+^CD4^+^ T cells (a), and splenocyte proliferation (b) and cytokine (c) recall response to MOG _35-55_ in mice treated daily with EcN or EcN^Lac^. **(d-g)** EAE development (d), IFN-γ^+^ and IL-17^+^CD4^+^ T cells in CNS (e) and spleen (f), and splenocyte proliferation recall response (g) to MOG _35-55_ in mice treated weekly with EcN or EcN^Lac^. **(h)** CFU in liver and colon from mice infected with *S. thyphimurium* after daily or weekly administration of EcN or EcN^Lac^. **(i-k)** S2W1 tetramer^+^ CD4 T cells in colon (i), liver (j) and representative S2W1 tetramer staining in liver (k) 14 days after *Salmonella thyphymurium* infection. **(l,m)** HIF-1α MFI in Neutrophils, monocytes and T cells (l), and number of HIF-1α+ DCs (m) in small intestine as a result of EcN or EcN^Lac^ treatment. Data shown as mean±SEM. **p<0.01, *p<0.05, ns: p>0.05.

